# The roles of microtubules and membrane tension in axonal beading, retraction, and atrophy

**DOI:** 10.1101/575258

**Authors:** Anagha Datar, Jaishabanu Ameeramja, Alka Bhat, Roli Srivastava, Roberto Bernal, Jacques Prost, Andrew Callan-Jones, Pramod A Pullarkat

## Abstract

Axonal beading—formation of a series of swellings along the axon—and retraction are commonly observed shape transformations that precede axonal atrophy in Alzheimer’s, Parkinson, and other neurodegenerative conditions. The mechanisms driving these morphological transformations are poorly understood. Here we report controlled experiments which can induce either beading or retraction and follow the time evolution of these responses. By making quantitative analysis of the shape modes under different conditions, measurement of membrane tension, and using theoretical considerations, we argue that membrane tension is the main driving force that pushes cytosol out of the axon when microtubules are degraded, causing axonal thinning. Under pharmacological perturbation, atrophy is always retrograde and this is set by a gradient in the microtubule stability. The nature of microtubule depolymerization dictates the type of shape transformation vis à vis beading or retraction. Elucidating the mechanisms of these shape transformations will facilitate development of strategies to prevent or arrest axonal atrophy due to neurodegenerative conditions.

## Introduction

Neuronal cells do not divide and are rarely replenished when lost. As a consequence, they have to last an entire lifetime while maintaining their tubular extensions, namely dendrites and axons. This makes the nervous system particularly susceptible to degeneration causing debilitating conditions. Axons, owing to their extreme lengths, are particularly vulnerable. Abnormal morphological transformations like the appearance of varicosities along the axon or “beading” is a hallmark of a very wide range of neurodegenerative conditions of the peripheral as well as the central nervous system. Beading occurs under surprisingly diverse conditions (1) such as Alzheimer’s (2–5), Parkinson (6, 7), autoimmune encephalomyelitis & multiple sclerosis (8), ageing (9–11), ischemia and resulting hypoxia or inflammation (12–15), or in response to traumatic stretch injuries (16–20). Axonal vericosities are also observed in normal axons as en passant synapses or presynaptic boutons (21, 22).

Two distinct physical mechanisms have been proposed for axonal beading. In induced Alzheimer’s-like conditions or under other biochemical perturbations, organelles are found to accumulate in the swellings (2, 23). In cases where traumatic injury is mimicked by applying sudden stretch, microtubule splaying or breakage within the swellings has been reported (16, 18, 20). These observations led to a “traffic jam” hypothesis which suggests that the beads are the result of local accumulation of organelles resulting from defects or breakage along the microtubule tracks (2). A different mechanism has been proposed for beading resulting from stretch injury of nerves or sudden changes in external osmolarity that could occur, for example, as a result of inflammation (24–26). In this model increased membrane tension drives a periodic modulation of the axonal diameter by a mechanism akin to that previously studied in synthetic lipid membrane tubes and cell protrusions (27, 28). In the case of axons under hypo-osmotic stress, theoretical analysis points towards an interplay between cytoskeletal elastic stresses and elevated membrane tension which leads to a tubular-to-peristaltic shape transition, which agrees quantitatively with those experiments (25).

Axonal retraction, on the other hand, occurs extensively as a natural process during development, a process known as pruning, apart from being induced by injury (Wallerian degeneration) or diseases like peripheral neuropathies (29–31). It is also an essential process in maintaining neural plasticity in the brain. Axonal atrophy due to neurodegenerative conditions were initially thought to occur via damage to the soma (30). Recent studies, especially those using chemical isolation of the cell body, have, however, revealed a “dying-back” process initiating at the axon’s distal end (29–31). Microtubule breakdown occurs early during this process (17). The interplay between cortical F-actin and microtubules, axon-substrate interactions, or motor activity dictate the dynamics of axonal retraction (32–34). In addition, axonal degradation via retraction or beading can also be triggered by axonal crush or transection due to injury, a process known as Wallerian degeneration. Extensive degradation of microtubules, possibly due to elevated Ca^++^ triggered by transection, is observed in these degenerating axons (17).

Recent studies have revealed that shape transformations occur early in diseased conditions and are not signatures of apotosis (30). Despite a remarkable range of conditions leading to axonal atrophy via these shape modulations, the biomechanical factors driving axonal loss remain largely unknown. Although the molecular pathways responsible for neurodegeneration are diverse, cytoskeletal disruption, especially that of microtubules, occur early and is responsible for the eventual loss of the axon through beading or retraction. Hence, understanding how cytoskeletal integrity influence axonal stability is critical in devising strategies to prevent, arrest or even reverse neurodegenearation (29–31).

In this article, we report a time-resolved investigation aimed at understanding the biomechanical responses of axons that lead to axonal atrophy when the cytoskeleton is perturbed using either pharmacological agents or laser ablation. We first show that destabilized axons exhibit two distinct shape modes. Direct disruption of microtubules using Nocodazole results in a peristaltic shape mode or beading characterised by the formation of axonal swellings arising progressively from the distal to the proximal side. The swellings drift in a retrograde direction on the average, and coalesce over time leading to eventual axonal atrophy. Perturbation of F-actin, on the other hand, leads to a retracting front, again initiated at the growth-cone, but characterised by a single sharp variation in calibre. In both cases, the outer membrane tube remains intact over the entire length of the axon and only varies in caliber. By combining experimental data and theoretical considerations, we argue that these shape evolutions are driven by membrane tension. Further, we show that gradients in microtubule stability is responsible for setting the retrograde directionality of atrophy, and the nature of degradation of the microtubule core dictates whether atrophy occurs via beading or retraction. This study arrives at some general principles that dictate axonal shape stability and provide a framework to eventually connect molecular level process to axonal biomechanical responses that drive axonal atrophy.

## Results

### I. Investigation of axonal beading caused by microtubule depolymerization

Typical *in vitro* cultures of chick DRG neurons show axonal growth of 100 to 150 µm overnight. When subjected to the drug Nocodazole (Noco), which disrupts microtubules, these axons develop a modulated shape with a series of bulges separated by thinned segments as shown in **Fig. 1a**. This morphology is termed axonal *beading.* We observe that the process of shape change is always initiated near the growth cone and proceeds towards the soma (**Fig. 1a**). Beads move along the axon bi-directionally at short timescales (tens of seconds), but there is a net drift towards the soma at the timescale of several minutes (**Fig. 1a**). This movement of beads leads to coarsening via bead coalescence. At much longer times, of the order of 30-60 min., the net retrograde migration of beads results in complete axonal atrophy, leaving behind a trail of a thin membrane tube as can be seen better in **Movies-1**,**2**. Fluorescence microscopy images revealing how cytoskeletal components are distributed in the beaded axons are shown in **Fig. 1b**.

**Figure 1:**
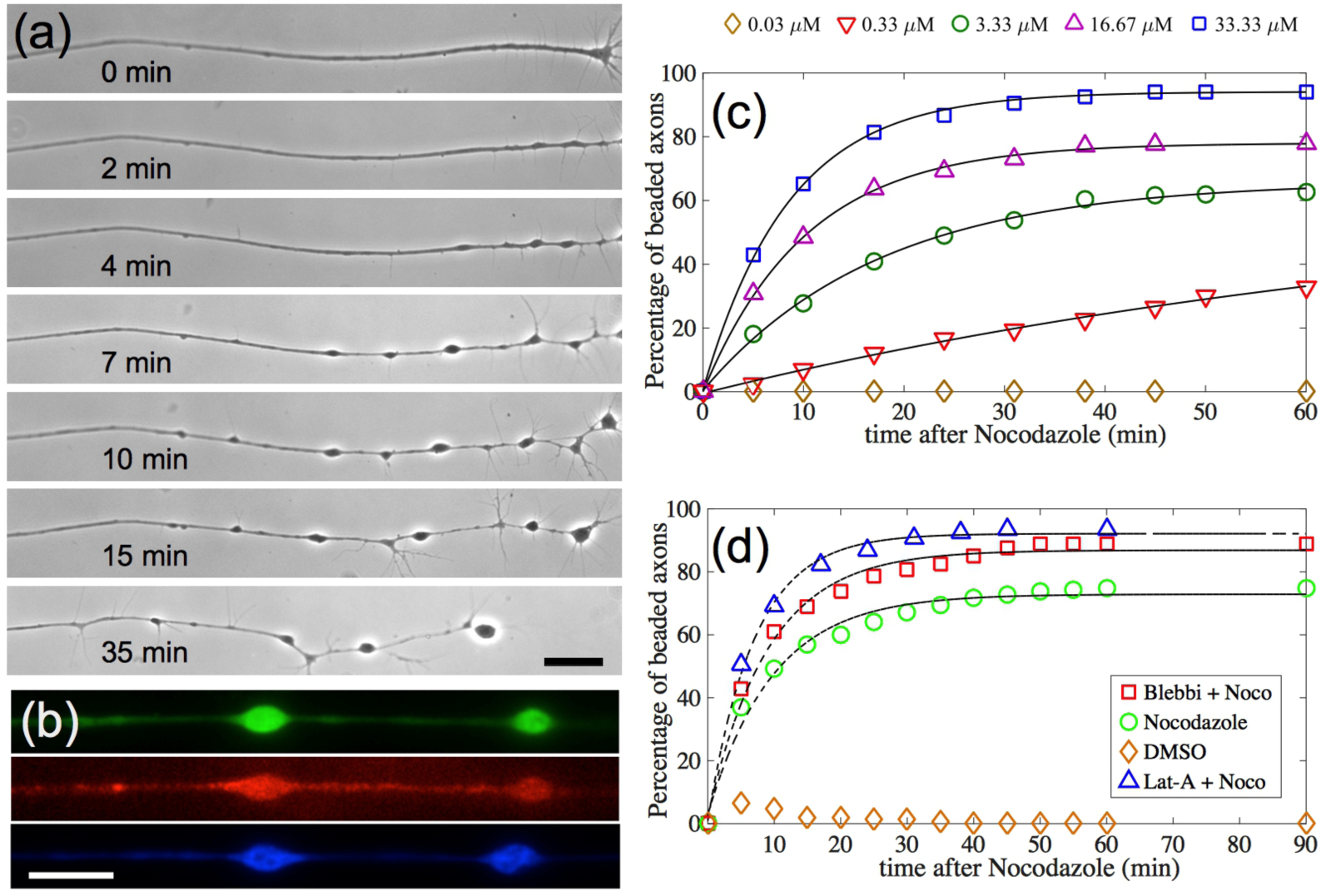
**(a)** Progression of Nocodazole induced beading in an axon. The time after addition of Nocodazole is indicated and the scale bar is 20 µm. Note that beading starts from the growth cone end (right) and progresses towards the soma (outside the field of view). This progression happens via retrograde bead migration, coarsening due to merging of beads, and formation of new beads. A thin membrane tube is left behind as beads migrate and coarsen (seen better in **Movies-1,2**). Further quantification of the bead distribution is provided in **Fig. S1**. **(b)** Fluorescence images showing the distribution of tubulin (green), F-actin (red) and neurofilaments (blue) in beaded axons (scale 20 µm). **(c)** Percentage of beaded axons over time for different Nocodazole concentrations. About 150 axons were recorded for each concentration. The black lines are fits to the function 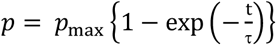. This yields a parameter τ, the beading time, which characterises the axonal susceptibility to beading. **(d)** Percentage of beaded axons for combinatorial drug treatments. Latrunculin-A was used at 1 µM, Blebbistatin at 30 µM and these pretreatments were followed by Nocodazole at 16.67 µM.

We then quantified the time evolution of Noco-induced axon beading, and what influence the acto-myosin cortex has on these dynamics. The percentage of beaded axons as a function of time for different drug concentration is shown in **Fig. 1c**. It can be seen from the exponential fits that the beading transition follows first-order kinetics with a threshold concentration between 0.03 and 0.3 µM. The choice of fit is justified by noting that the appearance of a bead following Noco treatment is expected to follow Poisson statistics. No beading is seen at 0.03 µM for over an hour of exposure time. The exponential dependence allows for a precise quantification of timescale for beading, which is a measure of the axonal stability to treatments, and are 83.4 min (0.33 µM), 17.6 min (3.33 µM), 10.3 min (16.67 µM) and 8.5 min (33.33 µM). An analysis of the relaxation times plotted as a function of Noco concentration yielding a measure of the rate of microtubule turnover in live axons as described in **Appendix 1 of Suppl. Mat**. The characterisation in **Fig. 1c** helps us to quantitatively investigate how disruption of F-actin or blocking of myosin-II affects beading. For this, axons were pre-treated with either 1 µM Latrunculin-A (Lat-A) for >10 min or 30 µM Blebbistatin (Blebbi) for >20 min and then subjected to 16.67 µM Noco. As can be seen from **Fig. 1d**, these pretreatments resulted in an increase in the rate of beading, with a characteristic time of 7.0 min for Lat-A and 9.1 min for Blebbi as compared to 10.3 min for control cells. Further quantification of the bead distribution is provided in **Fig. S1**. In order to crosscheck the Blebbi experiment, we pretreated cells with the Rho Kinase inhibitor Y27632 to reduce myosin light chain phosphorylation. This pretreatment did not have any noticeable effect on the total number of beaded axons (**Suppl. Mat., Table 1**).

**Table 1:** Quantification of beading or retraction events after pre-treating cells with the 30 µM Rho Kinase inhibitor Y-27632 for 20 min in each case. The quantification was done after 20 min of exposure to Noco at 16.6 µM or Lat-A at 1 µM.

| Drug | No of beaded axons | Total no of axons |
| --- | --- | --- |
| Nocodazole | 121 | 126 |
| Y-27632 +<br>Nocodazole | 123 | 143 |

**Table 1:** Quantification of beading or retraction events after pre-treating cells with the 30 $\mu\text{M}$ Rho Kinase inhibitor Y-27632 for 20 min in each case. The quantification was done after 20 min of exposure to Noco at 16.6 $\mu\text{M}$ or Lat-A at 1 $\mu\text{M}$ .
| Drug | No of retracted axons | Total no of axons |
| --- | --- | --- |
| Latrunculin A | 137 | 141 |
| Y-27632 +<br>Latrunculin A | 149 | 151 |

To understand why beading initiates from the growth cone side, we performed local application of Noco using a dual micropipette technique (see **Fig. S2**). No beading could be observed when the midsections or the proximal sections were exposed to the drug (**Fig. S2**). However, when growth-cones and the adjacent axonal shafts were exposed to Noco, beading started within a minute. This suggests that there is a growthcone-to-soma stability gradient for microtubules in the axon.

In order to test the prevailing hypothesis of axonal beading being caused by “traffic jams” in axonal transport (2, 20, 23), we recorded transport using both the phase-contrast method and using GFP-labelled synaptic vesicles (**Movies-3**,**4**). Although there is a tendency for organelles to pause within the swellings, several organelles can be seen passing through the beads unhindered during the early stages of Noco-induced beading as also shown in the kymograph **Fig. 2a**. As phase contrast images can only capture large and optically dense organelles like mitochondria, we also carried out imaging of tiny synaptic vesicles using synaptophysin-GFP. As can be seen from **Movie-4**, these vesicles too do not show any significant accumulation within beads. In both cases organelles occupied only a small fraction of the bead volume. Fluorescence imaging of microtubules in axons soon after beading clearly shows that continuous tracks of microtubules are left intact in most beads (n = 56 out of 75), as can be seen in **Figs. 2b,c,d**. It has been reported previously that post-translational modifications render additional stability to a fraction of microtubules in axons, which are depolymerised only upon prolonged exposure to Noco (35). Taken together, these observations suggest that accumulation of organelles is not the primary cause for Noco-induced beading, but rather a consequence of beading.

**Figure 2:**
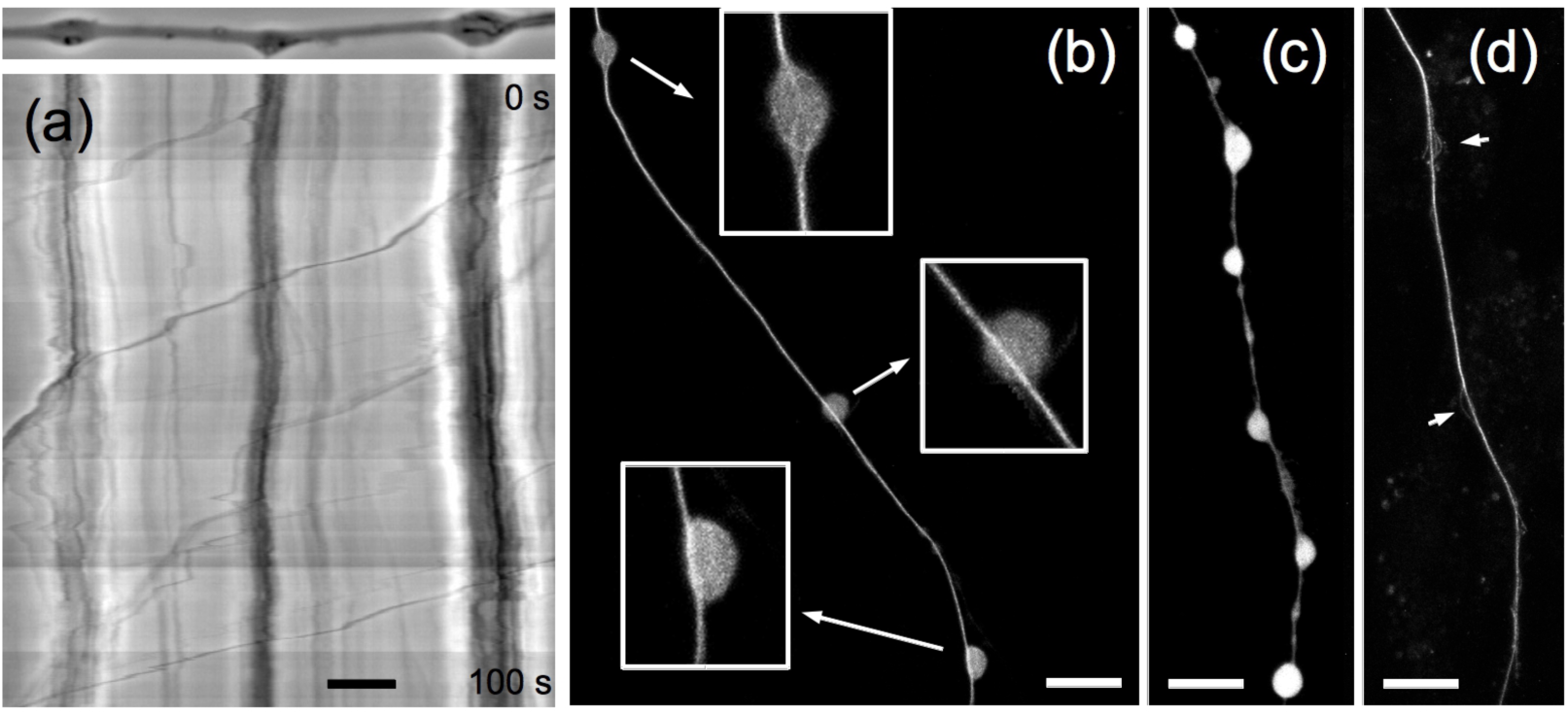
**(a)** Phase-contrast kymograph of a beaded axon showing frequent transport of organelles across beads with occasional pauses. The initial state of the axonal is shown on top. Time as indicated and scale bar is 10 µm. Also see **Movies-3,4**. **(b)** Cells with tubulin-GFP were fixed and free monomers were removed by mild Saponin treatment to reveal inner details (see Mat. & Mtd.). Contiguous microtubule (MT) bundles are often visible across beads (enlarged in lower insets) (n = 56 beads out of 75). A bead with disrupted MTs is also shown (top inset). **(c, d)** Continuous microtubule tracks in beaded axons is also seen after a complete removal of the membrane using Saponin+Taxol and then fixing. Images c & d were taken before and after this procedure, respectively. Arrows indicate splaying of microtubules seen in some beads. Scale bars: 10 µm.

We next looked at whether Noco treatment increases membrane tension, and thus would be the reason for beading (see **Suppl. Mat., Appendix-2** for reasoning). Indeed, a rapid increase in axonal membrane tension in conditions of hypo-osmotic shock has been shown to trigger axonal beading (25, 26). In Noco treated axons, such an increase in the membrane tension may occur due to generation of tubulin subunits leading to osmotic stress. The method of pulling tethers from axonal membranes allows us to estimate the membrane tension for Noco-treated and beaded axons (n = 12) to be 7.7 ± µNm^−1^ in comparison with control (n = 15) 5.6 ± 1.8 µNm^−1^ (see **Fig. S3** for data). In contrast, the critical membrane tension required to cause beading of an axon with unperturbed cytoskeleton is estimated to be of the order of 3×10^−4^ N/m (25). Thus, the small rise in membrane tension caused by Noco treatment alone is insufficient to cause axonal beading. However, as will be elaborated later, microtubule disruption drastically lowers the threshold tension due to a reduction of the bulk elastic modulus.

### II. Actin disruption causes microtubule dependent axonal retraction

To see how F-actin disruption affects axon response, cells were treated with 1 µM of Latrunculin-A (Lat-A). In this case, axons exhibit dynamics very different from Noco-induced beading as can be seen from **Fig. 3a**. Initially the growth cone collapses and its contents gradually accumulate and develop into a retracting front characterized by an abrupt change in radius. This single front propagates towards the soma, leaving behind a thin trail of membrane (**Fig. 3a**, **Movies-5,6**). In this respect this dynamics may differ from the usual axon retraction where the growth cone detaches and retreats. Occasionally, small beads appear close to the retracting front but these quickly merge with the front (**Movies-5,6**). The retraction speed, as can be deduced from **Fig. 3b**, remains almost constant till the entire axons becomes atrophied. The relative motion of the front and the extraneous particle seen in **Fig. 3a** suggests an extended flow field with velocity decreasing away from the front. Fluorescence microscopy images of F-actin, microtubules and neurofilaments in partially retracted axons show that all three cytoskeletal components are depleted in the thinned out segment of the axon (**Figs. 3c,d**). This shows that either cytoskeletal filaments are depolymerized or is being squeezed back along with the retraction.

**Figure 3:**
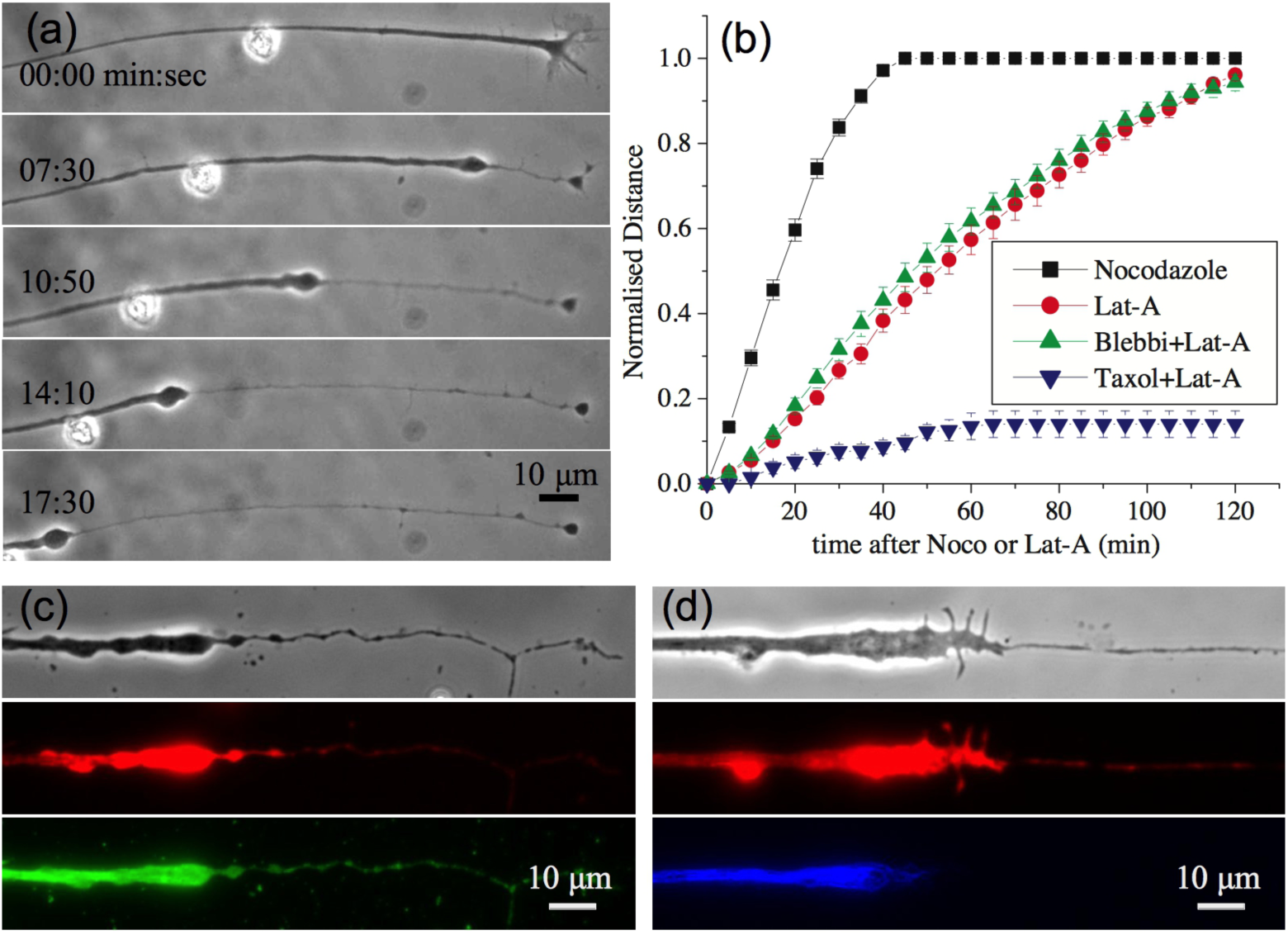
**(a)** Sequence of images showing axonal retraction in response to treatment with F-actin disrupting drug Latrunculin-A at 1 µM. The retraction front characterized by a sharp change in radius propagates towards the soma leaving a thin tube behind. Note that frames 2-5 have the same time gap. The relative movement of the front and the extraneous particle indicate a flow profile with decreasing velocity away from the front (also see **Movies-5,6**). **(b)** Plots showing the retracted length (thinned segment) measured from the growth cone end and normalised with total axon length for control axons; those pre-treated with the myosin-II inhibitor Blebbistatin; and those pre-treated with the microtubule stabilizer Taxol. The length of the axonal segment over which beading can be seen after treatment with Noco, also measured from the growth cone, is shown for comparison. Each curve is an average taken over n > 20 axons and the error bars are standard errors of the mean. The non-normalized and non-averaged data for Lat-A and Noco are shown in **Fig. S4**. **(c, d)** Phase contrast (greyscale) and corresponding fluorescence images showing F-actin (red) and microtubule (green) on left side and F-actin (red) and neurofilament

In order to check if microtubules are being depolymerized during retraction, neurons were pre-treated with the microtubule-stabilizing drug Taxol. Surprisingly, when axons (n=27) were treated with Taxol at 10 µM for 15 min prior to addition of Lat-A, it prevented axonal retraction except for a collapse of the growth cone (**Fig. 3b**, **Movie-7**). This suggests that Lat-A treatment also affects the microtubule stability in axons via an unknown signaling mechanism (36, 37). Retraction involves microtubule disassembly rather than intact filaments being squeezed out along with the front, unlike in retraction induced by nitric oxide (33). F-actin filaments, however, do not undergo complete disassembly and may be partly squeezed back as can be seen from rhodamin-phalloidin labelling, which marks only the polymer form of the protein (**Figs. 3c,d**).

We next explored the influence of myosin-II in Lat-A-induced axon retraction. We found that the retraction rate is slightly enhanced by treatment with Blebbi (**Fig. 3b**). Reducing myosin light chain phosphorylation by pre-treating cells with Y27632 also did not affect the occurrence of retraction events (**Suppl. Mat., Table 1**). Thus, as in the case of beading, myosin-II activity does not play a major role in this type of axon retraction, although contractility has been implicated in spontaneous axonal shortening (38).

Noco and Lat-A induced morphological transitions, vis à vis beading and retraction, show interesting concentration dependences. When neurons were treated with a much higher dose of Lat-A (10 µM), axons started to show more frequent beaded structures near the retraction front (**Table 2**). In some rare cases beading was extensive (**Fig. S5**). Conversely, when cells were treated with very low concentrations of Noco, a small fraction of axons showed retraction, without clear beading (**Table 2**, **Fig. S5**). Taken together with the fact that Taxol arrests Lat-A-induced retraction, these results suggest a common microtubule dependent mechanism for both retraction and beading, and this will be elaborated in the Discussion.

**Table 2.** Quantification of the concentration dependence of percentage of beading and retraction upon different drug treatments. Note that while Nocodazole predominantly causes beading, some percentage of retraction is also observed, and this number is higher at lower Noco concentration. And in the case of Latrunculin-A, the predominant mode is retraction, but an increasing percentage of beading close to the retracting front is observed with increasing Lat-A concentration.

| Drug | Concentration | Beading (%) | Retraction (%) | No. of axons |
| --- | --- | --- | --- | --- |
| Nocodazole | 3.33 $\mu\text{M}$ | 71.26 | 12.64 | 90 |
| Nocodazole | 33.3 $\mu\text{M}$ | 95.2 | 4.81 | 104 |
| Latrunculin A | 1 $\mu\text{M}$ | 3 | 97 | 100 |
| Latrunculin A | 10 $\mu\text{M}$ | 18.75 | 81.25 | 75 |

### III. Laser ablation causes bi-directional beading or retraction of axons

As mentioned earlier, biochemical perturbations always trigger shape changes in axons from the growth cone end. As shown by local drug treatment (see **Fig. S2**), this breaking of symmetry arises due to a gradient in cytoskeletal stability. Since axonal microtubules are arranged in a polar fashion, a directionality arising from axonal polarity need also to be investigated. Since local application of drugs to mid sections did not evoke any response, we resorted to point laser ablation. When ablated, most axons undergo complete transection and buckling (**Figs. S6a,b**). Every transected axon exhibited a rapid (< 1 s) shortening of length, which we call “snapping”, followed by a slower retraction. The snapping response, quantified in **Fig. S7**, and buckling show that axons are under tension before ablation. Interestingly, a subset of ablated axons does not undergo complete transection, presumably due to a lower laser power resulting from variations in focussing of the laser on the axon. In such “partial cuts” axonal segments on either side of the ablation point undergo thinning via either retraction or beading. This thinning response is initiated at the point of ablation and widens with time as shown in **Figs. 4a,c**, **Movies-8,9**).

**Figure 4:**
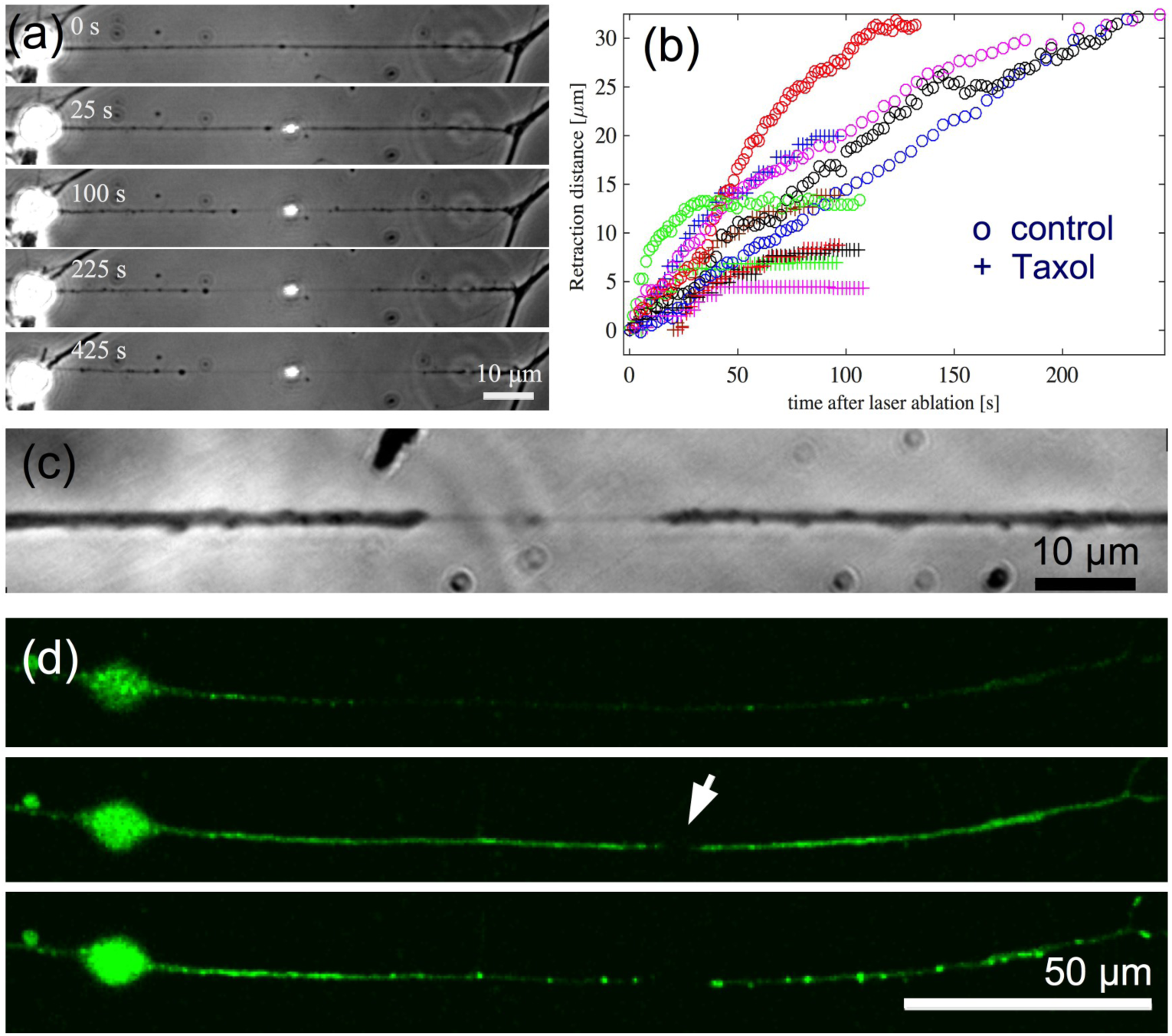
(a) Image sequence of an axon subjected to a single-shot “partial laser ablation” which leaves the membrane tube intact. The membrane tube starts thinning down starting from the point of ablation and this section widens with time. Beading is visible on the left side and retraction on the right in this example. The bright spot is due to the scattering of microscope light from the laser damage of the glass coverslip. (b) Plot of retraction distance measured from the point of cut as a function of time for control axons (open circles) and for axons treated with Taxol (+), obtained using partial cuts. (c) Image of another partially cut axon recorded at higher magnification shows the thin tube between the two retracting fronts. (d) Ca^++^ imaging in partially cut axons (n = 6 out of 6) shows an increase in Ca^++^ levels immediately after ablation (mid frame with arrow indicating the ablation point) and subsequent concentration of Ca^++^ in puncta along the axon, presumably due to sequestering of Ca^++^ into stores.

To proceed further, we defined a retraction front based on the change in axon calibre, as in the case of Lat-A induced retraction. This retraction behaviour for control axons is quantified in **Fig. 4b**. Most axons undergo complete retraction. Interestingly, pre-treatment with Taxol, which stabilizes microtubules, halts the retraction much earlier compared to untreated cells as is shown in **Fig. 4b**. This suggests that ablation affects microtubule stability in axons, presumably due to the creation of unprotected ends and membrane activated Ca^++^ influx as reported in literature (19, 39, 40). Indeed, fluorescence imaging of Ca^++^ show an marked elevation in Ca^++^ levels subsequent to ablation (**Fig. 4d**). These experiments show that even though axonal microtubules are arranged in a polar fashion with their plus ends pointed towards the growth cone, this polarity is not responsible for setting the propagation direction in axonal atrophy via beading or retraction. Instead microtubule stability gradients, in this case due to cut ends and Ca^++^ influx, dictate the direction of propagation of axonal atrophy.

### IV. The axonal actin-spectrin skeleton affects beading and retraction dynamics

Next we explore the effect of the recently discovered axonal actin-spectrin membrane skeleton on beading and retraction dynamics (41). This skeleton forms a 1-D periodic scaffold and is known to influence axonal caliber (42), circumferential tension (43), and longitudinal viscoelasticity (44). For this we perform spectrin knock-down experiments by treating the cells with beta-II spectrin specific Morpholino (SMO) for 48 hrs (see (44) for details). Axons thus treated were subjected to cytoskeletal perturbations using either 16.6 µM of Noco or 1 µM of Lat-A. For comparison, experiments were also done using a non-specific control morpholino (CMO). All knock-down experiments were performed on 2-DIV cells as the spectrin lattice was seen to be more prevalent in them (see (44) for quantification). The results obtained are shown in **Figs. 5a,b**.

**Figure 5:**
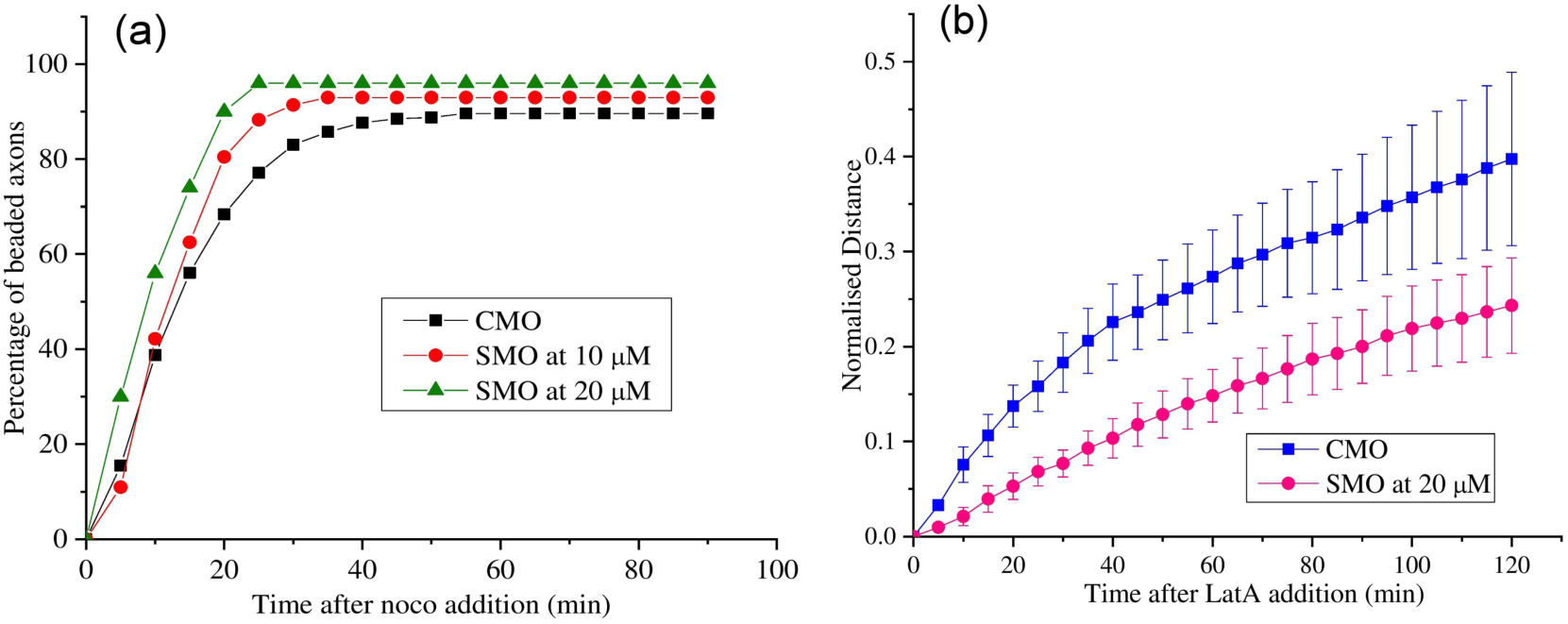
(a) Percentage of beading exhibited by neurons pretreated with either beta-II spectrin knock-down morpholino (n = 128) for 10 µM SMO and n = 50 for 20 µM SMO) or 20 µM control morpholino (n = 154) and then exposed to 16.6 µM Noco. (b) Averages of the retraction distances (length of the thin segments) normalised with the corresponding initial axon lengths for spectrin knock-down axons (n = 5) and control axons (n = 5) exposed to 1 µM Lat-A (error bars are SE).

It can be seen from **Fig. 5a** that axons with disrupted spectrin skeleton show a relatively higher percentage of beading for a given exposure time to Noco compared to control. The total number of beaded axons at saturation (long time) too increases with higher extent of knock-down. When spectrin knock-down treatment was applied to axons before treating with Lat-A, a marked reduction in the retraction speed is observed (**Fig. 5b**). These experiments clearly show that the axonal spectrin skeleton does play a role in axonal stability against beading and retraction induced by these drug treatments.

## Discussion

### Microtubule disassembly modifies axon shape

Axonal beading and retraction are hallmarks of a wide variety of neurodegenerative conditions that lead to axonal atrophy. Such commonality suggests a general mechanism behind these morphological changes. However, aside from the hypothesis that traffic jams of organelles are responsible for beading, there is no mechanistic understanding of such axonal shape modifications. Nonetheless, as cytoskeletal changes are known to be implicated in a variety of cell deformations, we inquired into how different types of perturbations to the axonal cytoskeleton might affect axon shape.

The three types of axonal shape change experiments performed in this study are summarized in **Fig. 6a**. These experiments – Noco-induced beading, Lat-A-induced retraction, and laser ablation – all involve modifications to the axonal cytoskeleton. Moreover, they all illuminate a key role played by microtubules in maintaining axon integrity, as revealed by Taxol pre-treatments. In the case of Noco treatment, which directly disassembles microtubules, loss of axon integrity occurs through a sequence of shape transformations involving the appearance of “beads” (or “pearls”). In contrast, when axons are treated with Lat-A, whose primary target is F-actin, they undergo a retraction process involving a front moving from the growth cone towards the soma that separates a thick, cytoskeleton-filled axon from a cytoskeleton-devoid membrane tube. Disassembly of microtubules following Lat-A is an indirect consequence, possibly arising from an interdependence between F-actin and microtubule mediated by other molecular players like tip-binding proteins, spectraplakins, etc (36, 45, 46). It is also known that freshly formed tyrosinated microtubules depolymerize within 10 min of treatment with Lat-A (37). Consistent with these results, retraction is significantly impeded by taxol treatment, which is known to stabilize microtubules against disassembly. Moreover, axon retraction following partial laser ablation can also be largely blocked by prior taxol treatment. Taken together, these experiments point to a key role played by microtubules in maintaining axon shape and calibre.

**Figure 6:**
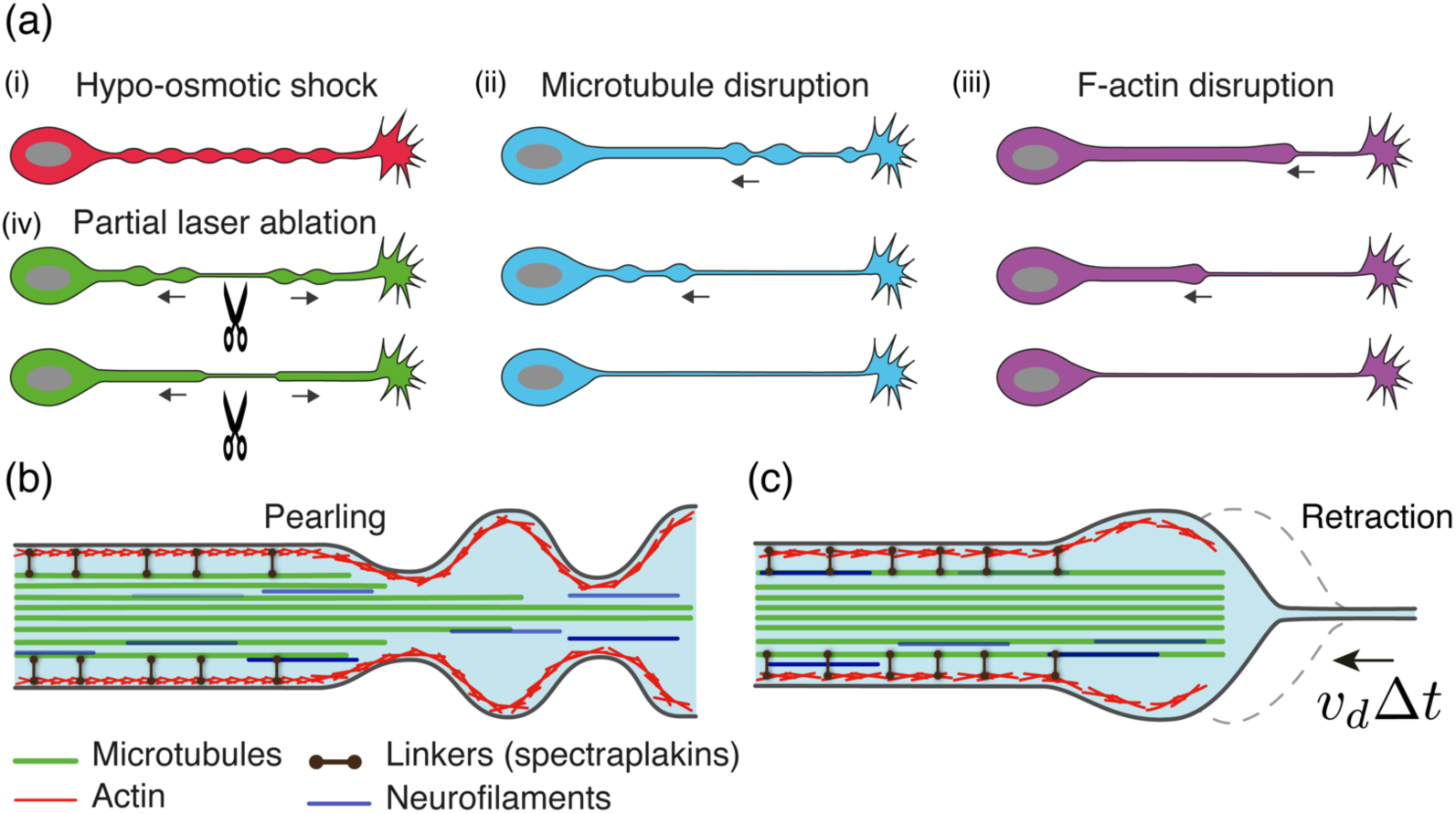
Shape transformations observed in axons under different induced conditions. (a) Axon beading or retraction occur following four types of perturbation: (i) hypo-osmotic shock produces a transient, periodic, non-propagating peristaltic mode (25). Moreover, this shape transformation occurs simultaneously along the entire axon and is transient (**Movie-10**). (ii) After microtubule disruption using Noco, beading always begins from the growth cone end and proceeds towards the soma. Finally, only a thin tube is left behind. (iii) In the case of F-actin disruption, a retracting front forms near the growth cone, and proceeds towards the soma, finally leaving a thin tube behind. (iv) In the case of partial laser ablation either beading or retraction occurs. In both cases, the ablation point thins down and expands with time, causing atrophy. Beading induced by Lat-A and laser ablation are prevented by taxol pre-treatment. (b) Experiments suggest that Noco treatment results in a radially shrinking microtubule bundle, separating it from the actin-spectrin-membrane skeleton. Membrane tension renders the resulting fluid contained between the membrane and remaining microtubule core susceptible to Rayleigh-Plateau instability. Axon beading thus occurs, which propagates from the thinner end of the bundle (near the growth cone), to the thicker one (near the soma). (c) Upon Lat-A treatment, axon cytoskeleton is disassembled in a front-like manner. This front separates a thin tube from a cytoskeleton-filled axon nearer to the soma.

### Microtubule stability gradients sets direction of beading and retraction

In both Noco-induced beading and Lat-A-induced retraction, axonal shape change is initiated near the growth cone and then proceeds towards the soma, accompanied by a flow of axoplasm and organelles (**Figs. 1a, 3a**). Is this retrograde directionality set by microtubule polarity or by microtubule stability? Local application of Noco using the dual micropipette arrangement shows that the cell shape is more readily affected at the growth cone end, when compared to the rest of the axon, showing the existence of a proximal to distal stability gradient (**Fig. S2**). These findings are consistent with studies reporting a correlation between microtubule stability and levels of post-translational modification or turnover rate, with less stable microtubules occurring closer to the growth cone (37, 47). They are also consistent with our observations that taxol treatment arrests Lat-A-induced retraction from the growth cone to the soma, indicating that microtubule stability gradient plays a role in setting the direction of front motion. Finally, upon partial ablation in the middle of the axon, retraction proceeds equally on both sides of the cut, and this retraction can be opposed by taxol treatment. In this case, additional stability gradients are generated by the ablation via the creation of a large number of unstable microtubule ends and an influx of Ca^++^ as shown in **Fig. 4d**, and reported by others (19, 39, 40). Taken together, these experiments show that microtubule stability gradients dictates the direction of propagation of beading or retraction.

### Axon beading is not a result of traffic jams

Based on the observation of accumulated organelles and hindered transport, it has been proposed that localized disruption of transport or “traffic jam” may be the cause for beading in some cases (2). This is an attractive model considering the fact that microtubule breaks or disorganization has been observed in axons with mechanically induced injury-like conditions (18–20). However, we note the following from our experiments. (i) Continuous microtubule tracks are seen in a large fraction of beads (n = 56 out of 75). (ii) Consistent with this, imaging of organelles show little hindrance at early stages of beading. No accumulation of synaptic vesicles is seen in beads at early stages. (iii) Bead locations are not fixed and beads migrate along the axon as they form. When beads move, fresh beads do not appear in place of the old ones, as is expected if microtubule defects dictate the location of beads. (iv) As will be elaborated below, the bead shapes—lemon-like at early stages and clam-shell-like at late stages—are typical of a surface maintained under tension as opposed to a shape defined by a collection of organelles. All these observations suggest that accumulation of organelles may not the primary cause for beading but rather a consequence of beading. Indeed, we observe splaying out of microtubule in beads which may be caused by the expansion of the surrounding cortex (**Fig. 2d**). In the next section we argue that beading and retraction can be explained via mechanical balance between membrane tension and cytoskeletal elasticity.

### Axon beading is a tension-driven shape instability

Cylindrical to beaded-like shape transitions, known in the physics community as “Pearling Instability”, have been studied previously in synthetic membrane tubes (48) and has been observed in *in-vitro* membrane tubes with a destabilized microtubule core (49). These systems lack molecular motors and beading is driven by membrane tension. Beading occurs also in cellular protrusions containing only F-actin (28), and in neurites subjected to osmotic shock (25, 50) (**Movie-10**). In these systems quantitative analysis have shown that beading occurs when membrane tension exceed a threshold value and the periodicity of beads increases linearly with radius as is expected for pearling instability. We observed that neurites of mouse PC12 cells exhibit beading when treated with either Noco or Lat-A. After Noco treatment, these neurites show a periodic shape modulation at early stages, and a linear relation between modulation wavelength and initial axon radius, as expected for pearling instability (25, 50) (**Fig. S8**, **Movie-11**). Thus, in all these cases, the beading process is related to the Rayleigh-Plateau instability of liquid jets (25, 50) (see **Appendix 2** in Supplementary Material for a simplified explanation).

As further evidence that axon beading is not the result of organelle “traffic jams”, but rather results from membrane tension, we analysed the bead shapes after Noco treatment. At early stages of beading lemon-shaped, axisymmetric beads are often seen (**Fig. 1b, S9**, Movie-1, Movie-2). At later stages, asymmetric, “clam-shell” beads are also observed (**Fig. 2b**). These shapes are strikingly similar to the quasi-stationary liquid beads, or pearls, that arise due to the Rayleigh-Plateau instability that dewets a liquid film initially coating a cylindrical fibre (51). In this case, the shape of the pearl derives from Laplace’s law, stating that the mean curvature of the drop is given by *H* = *P*/(2*σ*), where *P* is the internal pressure. At equilibrium, *P* is constant, and hence so is *H*.

We then inquired whether axon bead shapes could also be understood from a similar analysis. We first mathematically characterized the surface of a typical axon bead by determining the radial distance, *R*(*z*), from the bead centerline to the bead contour in the image plane as a function of distance along the axon, *z*; see **Fig. S9A**. Then, by fitting the contour data (**Fig. S9A**), we are able to calculate the bead curvature *H(z)*; see **Appendix 3** in the Suppl. Mat. for details. Interestingly, there is a central region of the bead where *H* is approximately constant, suggesting that the bead is liquid and with a surface shape governed by a competition between internal pressure and membrane tension. However, the mean curvature increases by roughly a factor of two near the bead edges (where the axon surface is expected to interact with the remaining microtubule core), a behavior which does not occur for liquid droplets wetting a fibre. This difference may be attributed to the bending stiffness of the axonal membrane, a property not present for a liquid droplet interface; see **Appendix 3** in Suppl. Mat.

We have seen here that Nocodazole drives a beading instability in axons, and an analysis of the bead shape and distribution suggest a tension driven process. Broadly, there are two ways that this could be triggered. First, Nocodazole increase the membrane tension, through osmotic effects arising from tubulin monomer generation, thus favoring beading. However, our optical tweezer measurements show no significant increase in membrane tension after Noco treatment (**Fig. S3**). This, then, points to the more likely scenario, which is that once microtubules are largely depolymerized, the elastic resistance to membrane tension-driven beading is lowered. More precisely, the critical tension for pearling instability is *σ*_*c*_ ∼ *ER*_0_ (25, 28); hence, a sharp reduction in the elastic modulus *E* caused by microtubule depolymerization, would lower the critical membrane tension, thus favoring beading.

### Beading vs. retraction

While Nocodazole treatment causes axon beading Lat-A treatment, in contrast, resulted in axon thinning via a retracting front. What might be the physical reason for this difference? Below, we argue that under the influence of an effective membrane tension, the shape mode that develops should depend on the nature of depolymerization dynamics of the cytoskeleton, and how these dynamics compare with those associated with the tension-driven Rayleigh instability.

- Based on **Figs. 2b,c,d**, that show a remnant microtubule trail after Noco treatment, we propose that Noco-induced disassembly occurs via radial thinning of the microtubule bundle. Since beads first appear near the growth cone, the radial thinning should occur fastest there and slowest near the soma, resulting in a shrinking conical microtubule core, with the point facing the growth cone. We furthermore assume that microtubule disassembly leaves behind in its wake a viscous fluid enclosed between the axonal membrane with its associated cortical skeleton and an intact microtubule core; see **Figs. 2b & 6b**. This fluid is assumed to contain cytosol, depolymerized cytoskeletal fragments, and neurofilaments (**Figs. 1b & 3d** show that neurofilaments are being squeezed into the beads or towards the soma respectively). As a result of membrane tension, this fluid is unstable to pearling.
- There is a typical speed associated with growth of axon beads. It is well-known that a liquid film of thickness *e* and viscosity *η* coating a fiber core of radius *r*_*a*_ – *e* is unstable for wavelengths *λ* > 2*πr*_*a*_, and these modes grow with speed 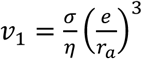 (51). This is the characteristic speed associated with flow of the cytosol. Furthermore, assuming a radially tapered microtubule core, and therefore a thickness of the fluid film that increases from the soma to the growth cone, beads will develop first near the growth cone and then progressively appear near the soma (52). This also accounts for the observation of bead drift (see **Movies 1,2, & 11**): because beads are more developed near the growth cone, and therefore the necks that connect them are thinner, the Laplace pressure increases from growth cone to soma, thus driving bead motion and coarsening (merging of fast moving beads nearer the growth cone with slower, adjacent ones). This situation is reminiscent of the dewetting of a radially tapered wire, as studied in Ref. (52). In contrast, under Lat-A treatment, experiments suggest that actin and microtubules depolymerize completely, starting at the growth cone, and hence no remaining microtubule core is left behind; see **Figs. 3c,d**. We therefore consider that Lat-A initiates a propagating depolymerization front with speed *ν*_*d*_ away from the growth cone. This depolymerization fluidizes actin and microtubules, and as a result, the depolymerized region of the axon is vulnerable to pearling instability. The velocity scale associated with the development of the instability depends on the membrane tension and the viscosity of the enclosed fluid (cytsol and fluidized cytoskeleton), and is given by 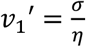. Assuming a bare membrane tension *σ* = × 10^−6^ N/m and *η*∼1 Pa.s (53), we find *ν*_1_′ ∼ 100 *μ*m/min, which is faster than the depolymerization speed implied by **Fig. 3b**, i.e., *ν*_*d*_∼ 100 *μ*m /100 min ∼ 1 *μ*m/min. This means that beading is happening very quickly compared with depolymerization, and in such a way that beads coalesce fast enough to give rise to a single retracting front. The enclosed volume is then expulsed towards the soma, forming a bulge just behind the depolymerization front; see **Fig. 6c**. We note further that the remaining thin tube does not undergo beading, because of the stabilizing effect of the membrane bending rigidity at this small radius (54).

### The role of the actin-spectrin membrane scaffold in axon shape stability

We observe that axons of spectrin knock-down neurons bead more easily compared to control and retracts slower (**Figs. 5a,b**). A passive spectrin skeleton is expected to stabilize axons against beading due to elastic effects (44). This is because the axonal spectrin tetramers are attached to the membrane by ankyrins and arranged longitudinally and hence is expected to get stretched during beading. This can explain the results shown in **Fig. 5a**. Recent experiments have shown that the actin rings that interconnect spectrin tetramers influence axonal caliber (42), and exerts a circumferential stress via myosin-II mediated contraction (43). It is conceivable that a circumferential stress arising out of the actin rings can contribute to the squeezing out of cytoplasm during axonal thinning. However, we do not observe any significant change in the retraction speed when myosin-II is pharmacologically inhibited (**Fig. 3b**). Therefore, understanding the precise biomechanics of the actin-spectrin skeleton during beading and retraction would require further experiments and theoretical analysis which invoke its anisotropic elasticity and active response.

### Conclusion

To conclude, we have shown that axonal beading and retraction are membrane tension driven instabilities occurring when microtubule integrity is compromised. The rate and manner in which microtubules disassemble—radial thinning with a spatial gradient vs. a decaying front—dictate the shape evolution, vis à vis beading or retraction. Moreover, we show that it is the microtubule stability gradient, and not polarity, that sets the retrograde flow of material during atrophy. These morphological transitions can be understood based on membrane and cytoskeletal mechanics. Furthermore, these shape changes bear similarities to inanimate systems, but with related underlying physics. For instance, both beading-like and retraction-like modes with directional propagation have been observed in the surface tension driven dewetting of tapered, liquid-coated wires (52).

Although microtubule stability has been long recognized as a major factor affecting neurons undergoing neurodegeneration, preventing or arresting axon loss using microtubule stabilizing drugs has met with limited success (55). While Taxol seems to prevent axonal loss in short term treatments like the one reported here, it is well known that prolonged anti-cancer treatment using Taxol and other microtubule stabilizing agents cause chemotherapy-induced peripheral neuropathy (56). Our study shows that axonal beading occurs even when a contiguous microtubule core remains. This is because the core becomes disconnected from the surrounding F-actin, and therefore also from the membrane, making the membrane unstable to beading. Moreover, we show that destabilization of F-actin triggers axonal atrophy by disrupting microtubules via unknown signaling pathways. Therefore, despite the central role played by microtubules in dictating the shape evolution, it is necessary to consider the stability of the microtubule-actin-membrane composite and not just microtubules in isolation. These interactions are mediated by a large number of other proteins (57). Thus, understanding the biomechanics of axonal shape evolution under conditions that perturb microtubules, as discussed in this article, provides a framework to connect diverse molecular level processes to general biomechanical principles that drive axonal atrophy.

## Materials and Methods

### Cell culture

Dorsal Root Ganglia (DRGs) from day-8 chick-embryo were incubated in 0.25% Trypsin-EDTA (Gibco 15400) for 10 min, dissociated in HBSS (without Ca^++^, Mg^++^), and plated on clean, uncoated glass coverslips. For plating, L-15 medium (Gibco 11415) made viscous using Methocel E4M (Colorcon ID 34516) at 0.3 gm to 50 ml, stirred overnight at 4 °C, and supplemented with 10% heat inactivated Fetal Bovine Serum (Gibco 10100), 33.3 µM D-Glucose (Sigma G6152), and Nerve Growth Factor NGF 7S (Invitrogen 13290-010) at 20 ng per ml was used. A good growth of axons (about 150 µm or longer) is seen after 14 hours of incubation at 37 °C. Before experiments, cells were incubated for 20 min in growth medium as above but lacking Methocel.

### Drugs

Nocodazole (Sigma M1404), Latrunculin A (Life Technologies L12370), Blebbistatin (Sigma-Aldrich B0560), and Taxol (Sigma T7402) were dissolved in DMSO with final DMSO kept below 1% v/v. DMSO control were performed at 1% DMSO. Y-27632 (Sigma Y0503) stock was prepared in water. All drugs were tested on chick fibroblasts for known responses.

### Immunolabeling and Ca^++^ imaging

Cells were fixed using 0.05% EM-grade glutaraldehyde (Electron Microscopy Sciences 16200), permeabilized using 0.5% Triton X-100 (Sigma 8787), and exposed to 5% heat inactivated goat serum (Abcam ab7481) for one hour at room temperature (all in PHEM buffer). After rinsing, the cells were exposed to the primary antibody, DM1A mouse monoclonal anti-α-tubulin (Sigma T6199) or mouse antigen antibody against neurofilament (Developmental Studies Hybridoma Bank, USA 3A10), at 1:1000 overnight at 4 °C. Cells were rinsed and exposed to the secondary antibody Alexa fluor 488 (Molecular Probes® A-11001) at 1:10000 for 1 hr in the dark. After rinsing the cells were imaged using Andor iXon 885 EMCCD camera at 100X magnification. For F-actin labelling, cells were permeabilized usind 1% Saponin, exposed to Rhodamine-Phalloidin (Fluka 77418) at 0.025 µg per ml for 20 min at room temperature, and then rinsed. Calcium levels in axons was imaged by preloading the cells with Fluo-4 AM (Invitrogen F14217) at 0.1 μM concentration. Images were taken using a Leica TCS SP8 confocal system before and after ablation with exactly the same imaging parameters.

### Visualization of microtubule tracks

Two techniques were used for this. (i) Cells expressing tubulin-GFP were treated with 10 μM Nocodazole for 5—7 min, fixed (as above) to retain the overall beaded shape and permeabilized using 1% Saponin for 1 min to get rid of free tubulin which otherwise causes strong background. (ii) After Nocodazole treatment as above, cells were treated first with a mixture of 1% Saponin and 10 μM Taxol for 1 min for complete removal of membrane and tubulin monomers without losing microtubules, and then fixed. In both cases images were acquired at 63X, 1.4 NA, using Z-stack feature of a Leica TCS SP8 confocal system.

### Tubulin-GFP

cells were obtained by transfecting cells with the plasmid using cuvette electroporation (NEPA21, Nepa Gene Co.). These cells were used after 48 hr of incubation. The construct was tested using chick fibroblasts and growth cones where individual microtubules are clearly visible.

### Laser ablation

Nd:YAG laser (Spitlight 600, Innolas, Munich, Germany) with wavelength 355 nm (UV) and pulse width 6 ns was used for ablation. The beam was directed into the back port of an inverted microscope (Olympus iX71) using steering mirrors and reflected by a dichroic mirror into the objective’s back aperture. Imaging was done using either a 40X (NA 0.7) or a 20X (NA 0.45) UV compatible Olympus objective and a CCD camera (Optikon, PCO). An infra-red lamp was used in order to keep the neuronal sample at 35 ± 2 °C as measured using an immersed tiny Platinum resistor. For Ca imaging, ablation was performed using a 355nm, 25 μJ, 350ps pulsed laser (TeemPhotonics, PowerChip PNV-M02510-100) coupled to a Leica TCS SP8 confocal microscope. Temperature was set to 37 ± 0.1 °C using the microscope incubation chamber.

### Image analysis

Kymograph was prepared using phase-contrast images recorded at 5 fps after addition of Noco at 16.6 µM and a custom program written in Matlab (MathWorks, USA). Shape analysis of the beads was performed using a custom edge detection algorithm written in Matlab. Edges were detected by finding the peaks in intensity gradients across the axon. The detected edge was fitted to a 7^th^ degree polynomial to get a smooth boundary before calculation of principle curvatures.

### Data availability

Data and protocols related to this article will be available from the corresponding authors on reasonable request.

## Acknowledgements

We thank Mohammad Ashraf Arsalan for the preparation of the Kymographs, Serene Rose David for help with experiments, Manoj Mathew and Sandhya Kaushaka for facilitating the usage of the laser ablation setup, Ashish Kumar Mishra for calcium imaging, Veena Chatti for help with transfection protocol, Thomas Bornschlögl and Patricia Bassereau for facilitating usage of the laser tweezer set up. ACJ would like to thank Adrien Daerr and Joshua McGraw for helpful discussions. RS was supported by a research grant from the Department of Biotechnology, Gov. of India (BT/PR13310/GBT/27/245/2009). PAP acknowledges the Department of Science and Technology, Gov. of India, for support via Ramanujan Fellowship.

## Supplementary Material

### Quantification of inter-bead distance and its dependence on initial radius

**Fig. S1:**
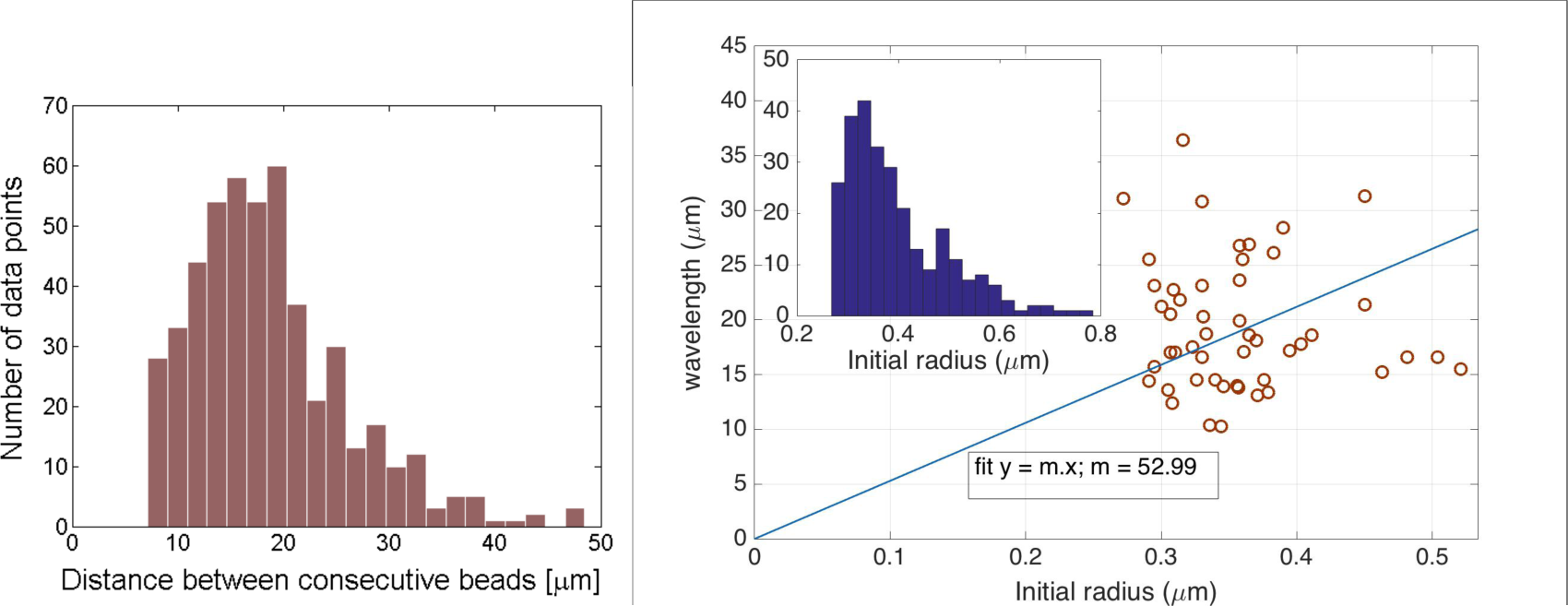
Characterization of beading as a result of 15 ± 3 min. exposure to 16.7 µM nocodazole. **Left:** distribution of distance between consecutive beads (n = 490 bead pairs). **Right:** plot of averaged distance between beads per axon (wavelength) versus the averaged initial radius of an axon (n = 50 axons). The fitted line was forced through the origin. This linear fit is motivated by the expected scaling of the most unstable beading wavelength with the initial radius of the axon, as predicted by Rayleigh-Plateau type instabilities (Ref. 49). The inset shows the distribution of initial radius (n = 272).

### Local application of nocodazole using dual micropipettes

**Fig. S2:**
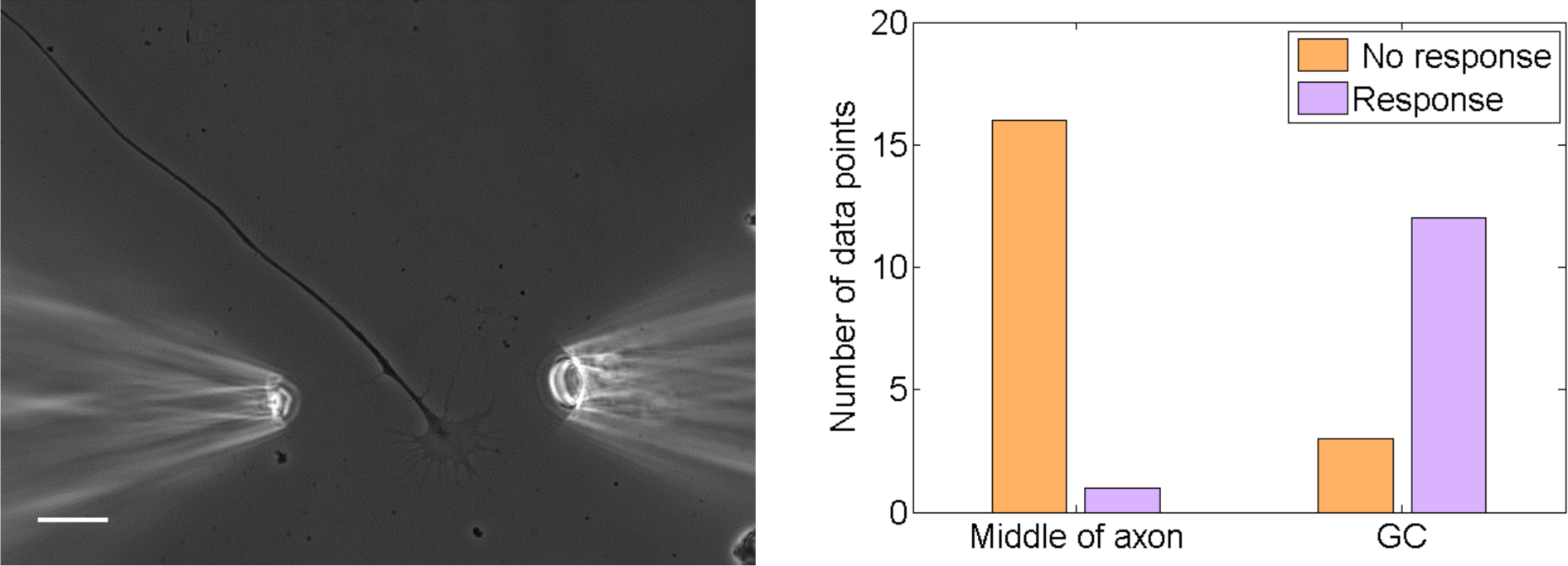
**Left:** Dual micropipette arrangement for local application of Noco at the growth cone (GC) of an axon, using an infusion pipette on left and a suction pipette on right (bar: 20µm). The infusion and suction are controlled using XenoWorks Digital Microinjector (Sutter Instruments Co.). The typical concentration profile around the application point was measured using a fluorescent dye instead of Noco. The drug exposure was restricted to about 10 µm of the axon shaft. **Right:** quantification of the beading response of axons to such local treatment, depending upon the location of drug application. Distal shaft next to the growth cone is seen to be much more vulnerable than the proximal shaft.

### Membrane tension measurements

**Fig. S3:**
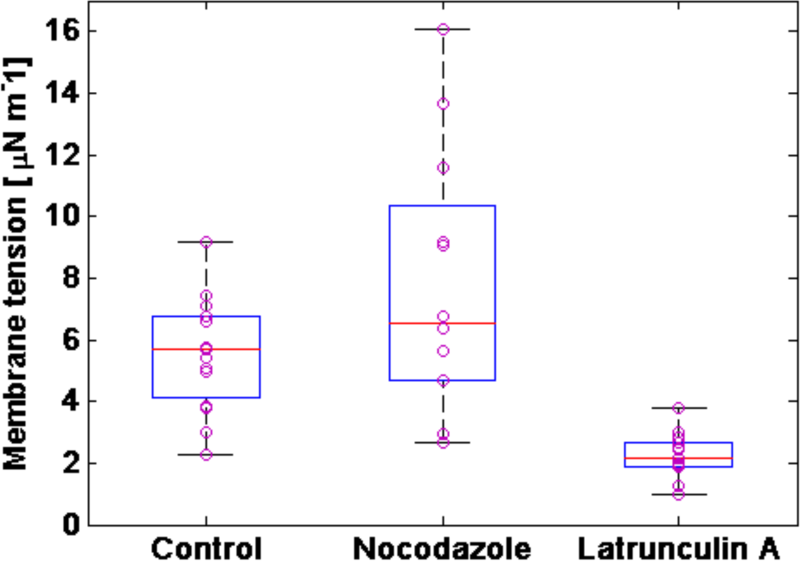
Membrane tension measured using steady state membrane tethers extracted out of axons using an optical tweezers (see Datar et al., Biophysical Journal, vol. 108, pages 489-497, year 2015 for details). Data for control cells, cells treated with 16.67 µM Nocodazole and cells treated with 1 µM Latrunculin-A are shown (n = 12 axons each). The horizontal red lines are the medians and the boxes are bound by the 25th and 75th percentiles in each case. The whiskers mark the extreme data points.

**Fig. S4:**
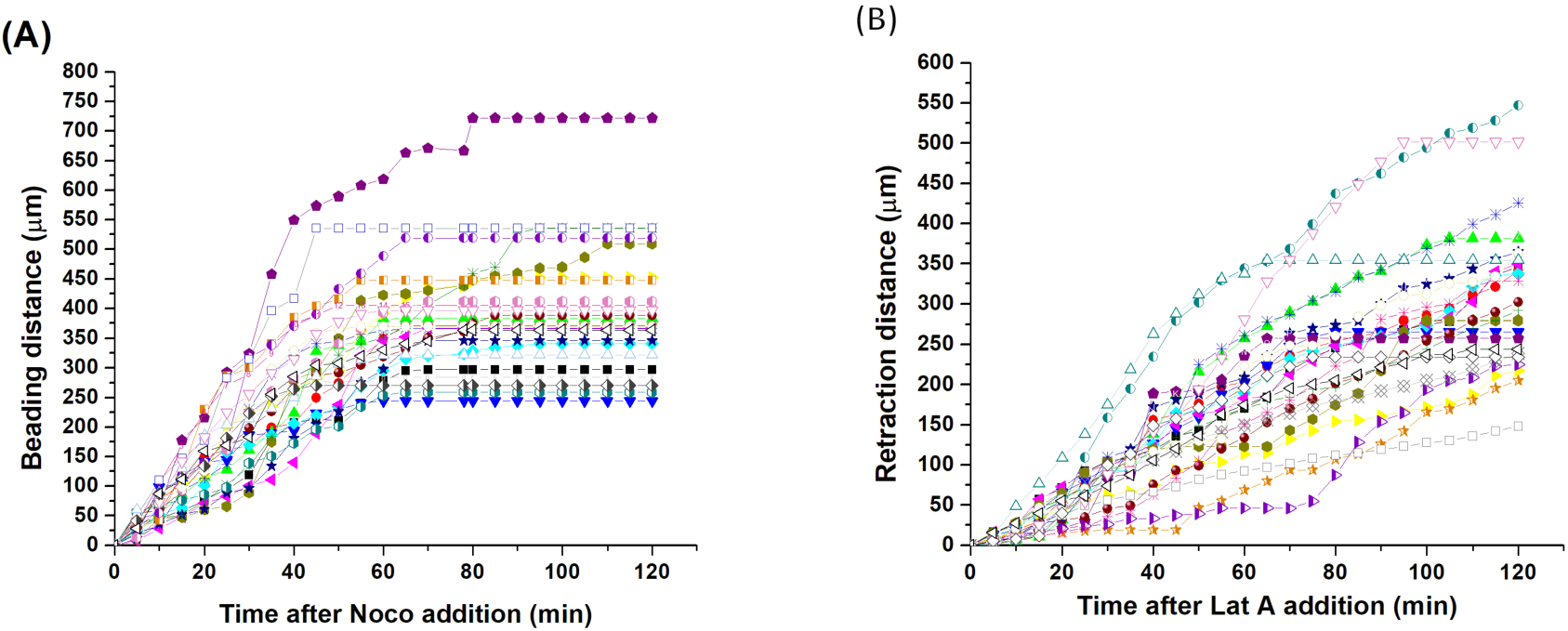
**(A)** Data for individual axons showing how the distance over which beading is observed grows with time. The distances corresponding to the last time point gives the length of the respective axon. Distances are measured from the growthcone. **(B)** Data for individual axons showing how the distance over which retraction has progressed grows with time. The distances corresponding to the last time point gives the length of the respective axon. The averaged data for both cases are shown in the main text.

**Fig. S5:**
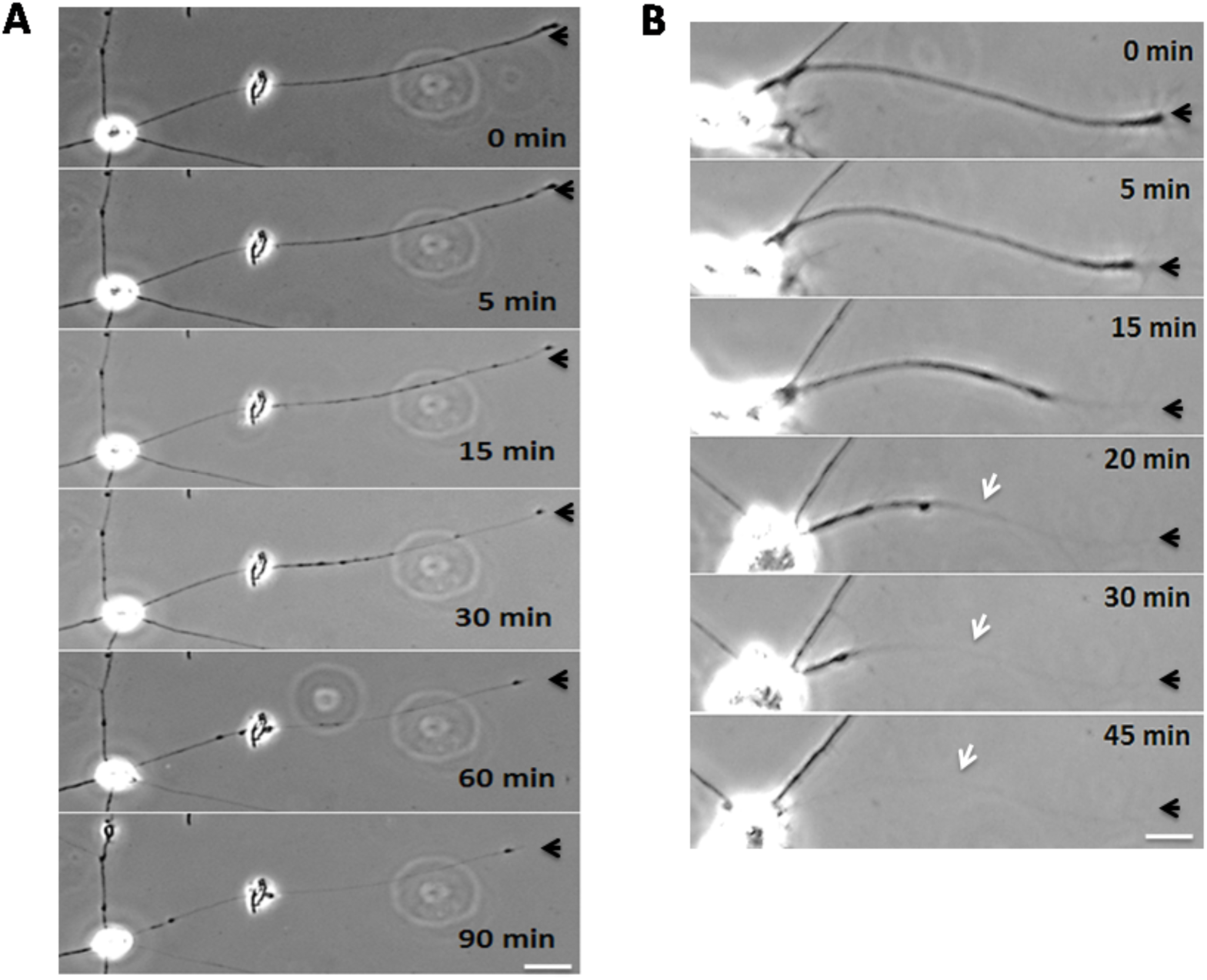
**(A)** Image sequence showing beading of axons treated with high concentration (10 µM) Latrunculin-A. **(B)** Image sequence showing an example of retraction happening under the influence of low concentration (3.33 µM) Nocodazole. Scale bars are 10 µm.

### Laser ablation

We observed that the immediate response of axons to laser ablation is of a very fast retraction away from the point of cut (**Fig. S5a**). We call it snapping. To test whether this is a mechanical response of the axon or just a laser induced damage, we performed the same experiment using cells fixed with glutaraldehyde. These cells did not show any significant snapping (**Fig. S5b**). Subsequent to the sudden shortening, untreated axons show buckling and/or beading and in some cases, folding back of the cut tips away from the point of laser ablation. Beading appears to progress away from the point of laser ablation (**Movies-8,9**).

**Fig. S6a:**
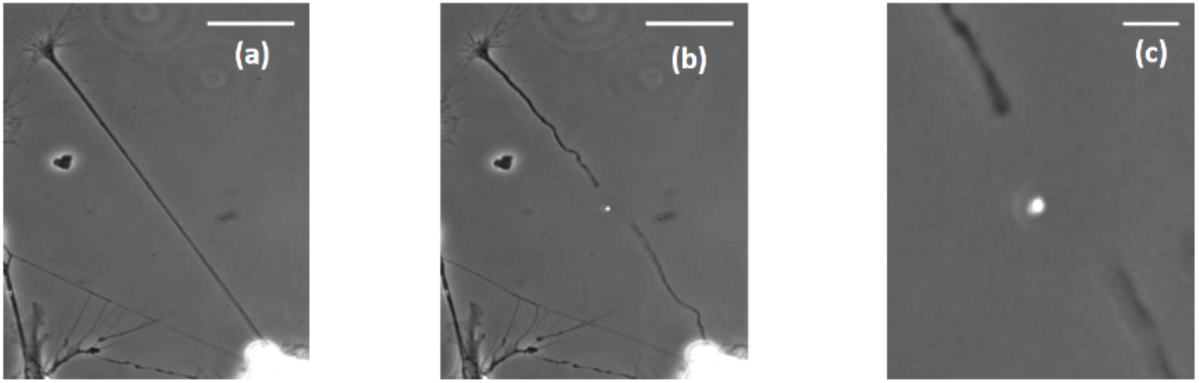
An axon before **(a)** and after **(b)** a laser cut (bar: 40 µm). Magnified part of image (b) is shown in **(c)** (bar: 8 µm). After the cut, axons under go a sudden retraction which accompanied by buckling and then a slower retraction.

**Fig. S6b:**
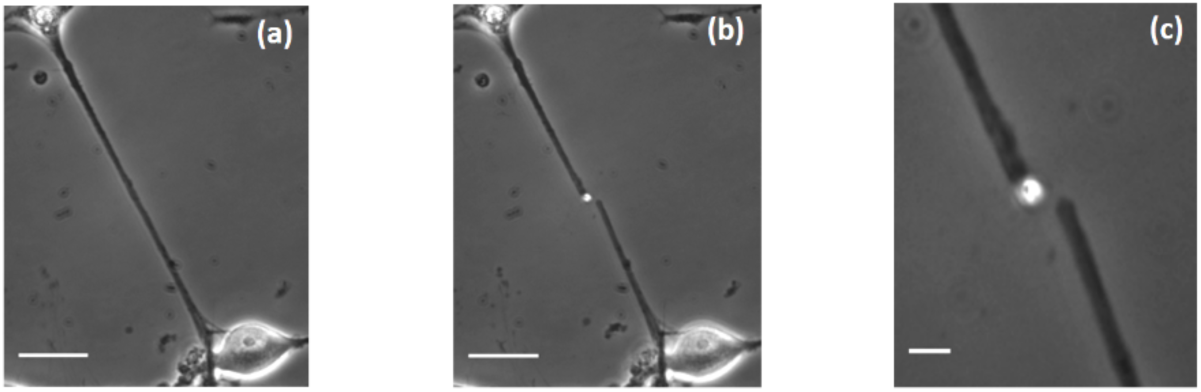
A glutaraldehyde-fixed axon before **(a)** and after undergoing laser cut **(b)** (bar: 40 µm). Magnified part of image (b) is shown in **(c)** (bar: 8 µm). Fixed axons do not show any significant shortening after the cut. This shows that the laser cut itself is clean and shortening in non-fixed cells is a true axonal response to transection.

### Distribution of snapping distance after laser transection of axons

**Fig. S7:**
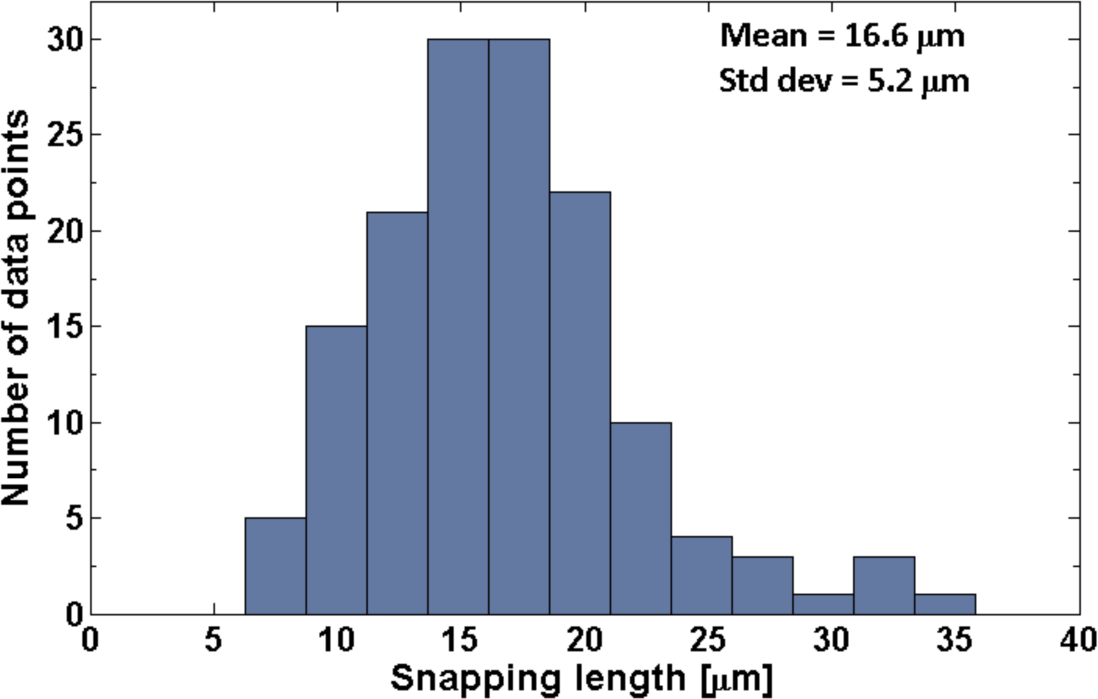
Distribution of snapping distance after laser transection of 72 axons, with mean 16.6 µm and standard deviation 5.2 µm. The snapping and buckling responses (see Fig. S6a) show that the axons are under significant rest tension.

### Beading observed in PC12 neurites treated with either Latrunculin-A or Nocodazole

**Fig. S8:**
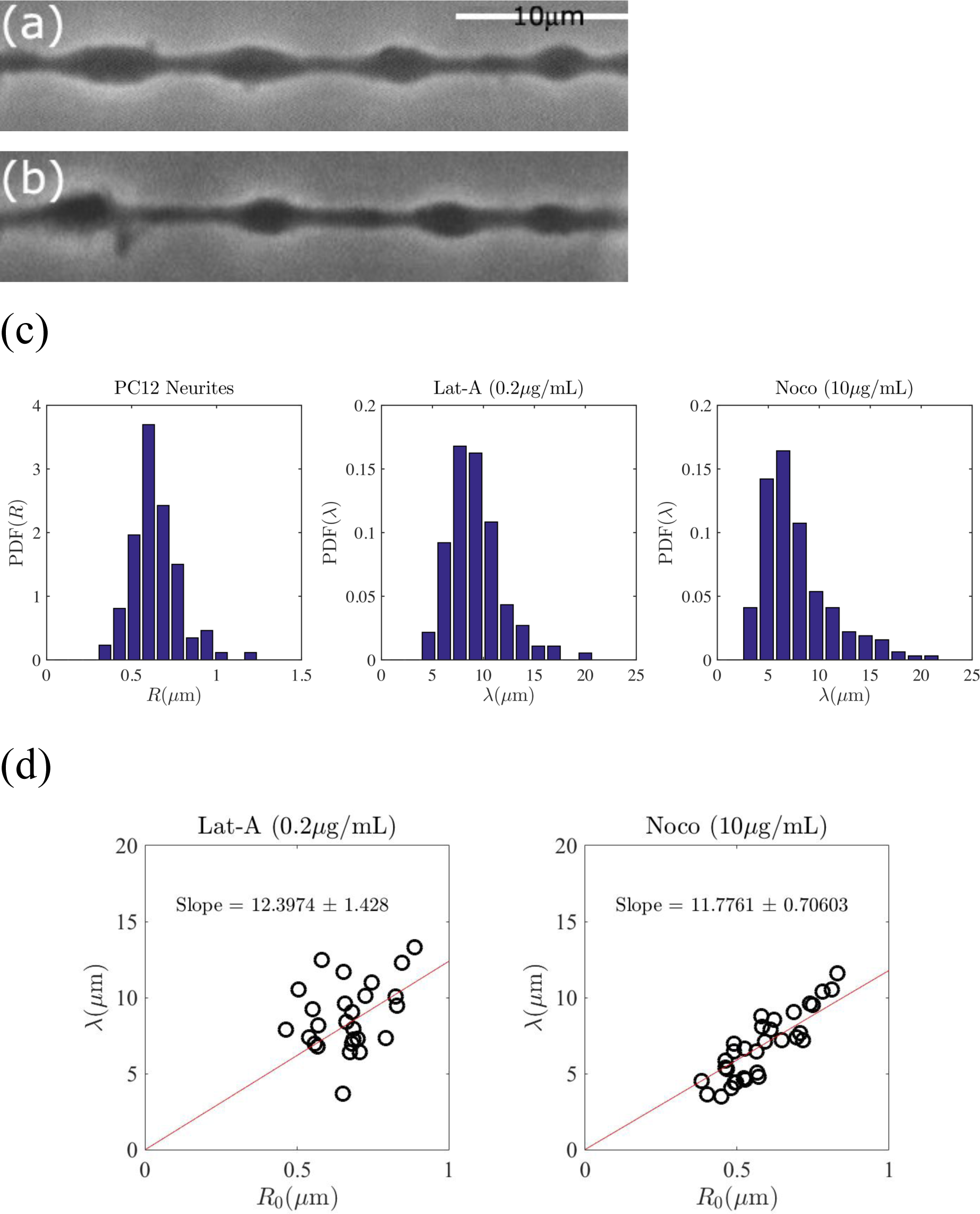
Neurites of mouse PC12 cells show beading when either microtubules or f-actin is pharmacologically disrupted. **(a)** Beading observed in PC12 cells when treated with 10 μg/ml Nocodazole (Noco). **(b)** Beading in PC12 cells occurring as a result of exposure to 0.2 μg/ml Latrunculin-A (Lat-A). **(c)** The distributions of the initial radius and the beading wavelengths for PC12 cells treated with either Noco or Lat-A as indicated on the plots. **(d)** Wavelength (λ) vs initial radius (R_0_) plots for PC12 cells treated with Noco or Lat-A as indicated above the figures. The fits are forced through (0,0).

### Analysis of bead shape

**Fig. S9:**
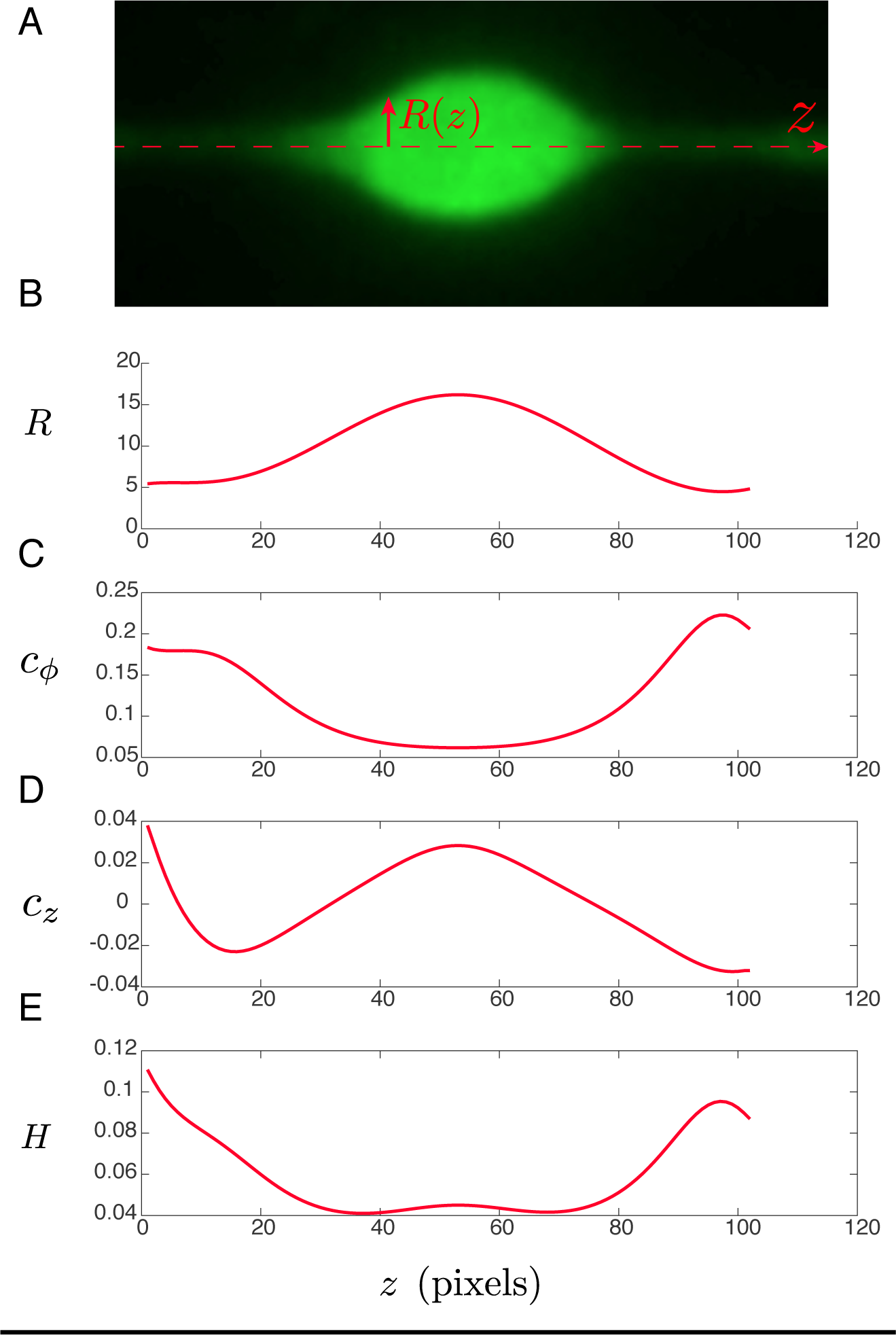
Analysis of bead shapes reveals balance of forces on axon surface. **(A)** Fluorescence image of axon bead after Noco treatment. **(B)** Radial distance from centerline to axon bead surface, *R*, as a function of axial position, *z. R* is in pixels. **(C)** Principal curvature of bead surface in plane perpendicular to *R* and *z*, as a function of *z*. **(D)** Principal curvature of bead surface in (*R, z*) plane, as a function of *z*. **(E)** Mean curvature of surface, *H*, as a function of *z*. Curvatures are expressed in inverse pixels.

## Appendix 1 Quantification of microtubule depolymerization dynamics

**Fig. S10:**
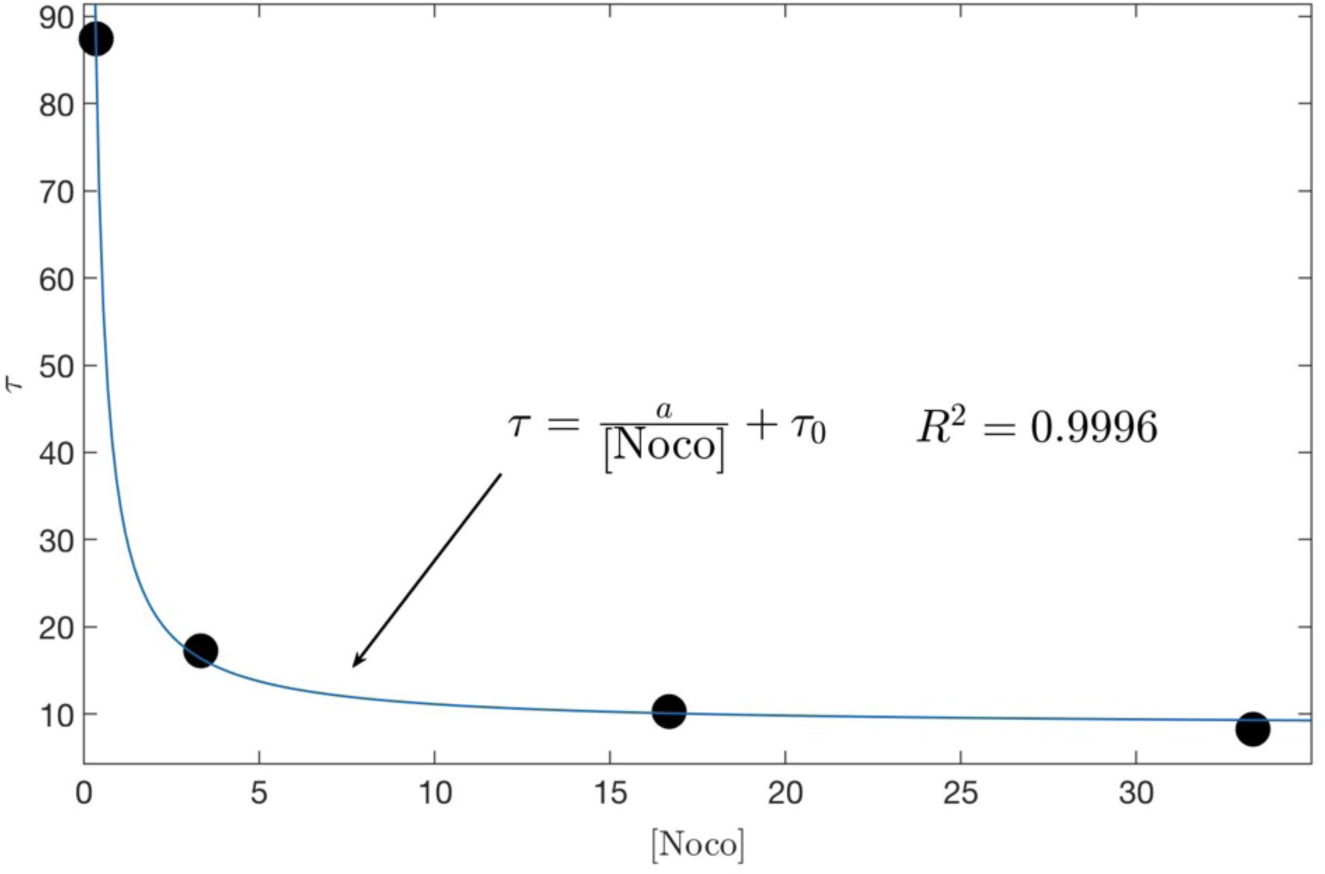
Plot of the beading time constant obtained from the fits shown in **Fig. 1c** plotted against Noco concentration [Noco]. The data is fitted to the equation shown in the figure and discussed in detail below.

The time constant *τ*([Noco]), corresponding to the rate of beading at different Noco concentrations and obtained from the fits shown in **Fig. 1c**, is plotted against Noco concentration in **Fig. S10**. This data fits well to the equation 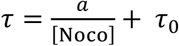, where *a* = 26 and *τ*_0_ = 8.5 *s* are constants. Saturation of the curve to *τ*_0_ indicates that there is a rate limiting step in the Noco-induced depolymerization process. The diffusion of Noco into the axon happens within a minute as per the response of the growth cone end of the axon observed in local drug experiments. This is fast compared to the time taken for beading which is of the order of 10 min at the highest Noco concentration (see **Fig. 1c)**. Therefore, we conclude that the rate limiting step has its origin in the turnover rate of microtubules in the axons. This is because Noco binds to tubulin in a 1:1 ratio, and at high concentration all *αβ*-subunits are bound to Noco and incapable in participating in the polymerization process. In such a scenario, the beading rate is dictated by microtubule depolymerization rate alone. This allows us to obtain an estimate for the microtubule turnover rate in live axons, which is otherwise difficult to quantify. If one supposes that the average length of microtubules is *L*∼5 μm (Yu and Baas, 1994), and that the bare shrinkage rate (*i.e*, without any Noco) is 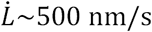 (REF), then *k*_off_ should be of order 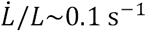. The inverse of this gives 10 s, which is close to the value of *τ*_0_ obtained from the fit in **Fig. S10**. Thus, quantifying the beading percentage under the action of Noco provides a powerful assay to quantify the stability of microtubules in axons. It can be used to study the effect of post-translational modifications to microtubules on their stability or to screen anticancer drugs for their potential to cause neuropathy.

## Appendix 2 Rayleigh-Plateau and Pearling instabilities

A cylindrical column of any fluid is unstable and will break up into a series of droplets. This break-up is driven by surface tension *σ*. This is because there is a net reduction in surface area in going from a cylinder of radius *R*_0_ to a series of *n* spherical drops of radius *R*_*s*_ of same net volume. Since the total volume has to be conserved, this gives a cylinder-to-spherical drops surface area ratio 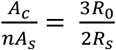. Thus, a series of spheres with 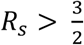 has a lower surface area compared to the original cylinder. As a result, a fluid cylinder is unstable to shape perturbations with wavelengths greater than *R*_0_.

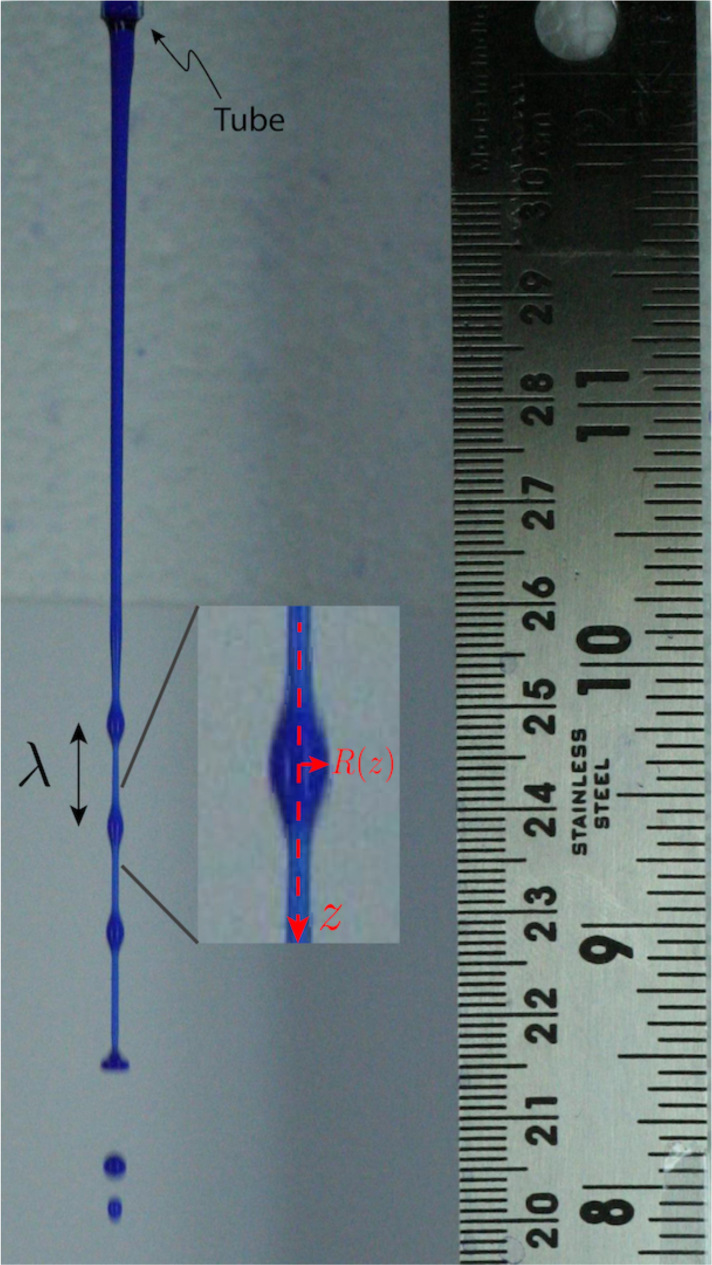

This is, of course, a static argument, and the actual shape change of the fluid cylinder depends on dynamics. It was shown by Rayleigh that the shape evolution happens via sinusoidal peristaltic modes. Sinusoidal perturbation of the radius of the form *R*(*z*) = *R*_0_ + *ε* sin(2*πz*/*λ*) reduces surface area provided the wavelength *λ* > 2*πR*_0_ (ignoring epsilon square terms which guaranty constant volume). In such cases, the higher Laplace pressure (*P* ∼ *σ*/*R*) in the thinner parts drive fluid to the thicker parts leading to the fomation of a series of bulges that grow with time until break-up. The optimal wavelength—the fastest growing mode—is decided by a competition between the gain in surface energy for a given wavelength and the time scale for fluid transfer. Long wavelengths (resulting in fewer final drops) are energetically more favourable but they are slow to grow as they require transport of fluid over long distances (trough to crest). Short wavelengths require fluid flow over short distances but also results in less energy gain and hence slow to grow. Thus, in experiments only the fastest growing mode becomes apparent.

An easy way to obtain a fluid cylinder is to allow fluid to fall free from a vertical faucet. The image on the left shows a snapshot of ink coloured water dropping from a plastic tube which can be seen at the top. The vertical axis can be thought of as time with zero time at the tube opening.

The mechanism is similar for a synthetic lipid bilayer membrane tubes filled with a fluid undergoing “Pearling instability”, expect that in this case the tube is unstable only if the membrane is under tension (an unconstrained piece of lipid bilayer has zero tension and hence zero interfacial elastic free energy). Moreover, the gain in surface energy has to overcome the cost of bending the membrane. Hence there is a small but finite critical tension for the tube to become unstable to Rayleigh-Plateau-like modes (see Bar-Ziv and Moses, *Phys. Rev. Lett.*, **73**(10), p.1392-1395, (1994)).

If the membrane tube is filled with an elastic matrix, like in the case of axons, there is a bulk elastic free enrgy cost for shape change as any deformation increases the free energy. This increases the threshold tension required for beading. In a highly simplified case, this critical tension goes as *σ*_*c*_ ∼ *ER*_0_, where *E* is the bulk elastic shear modulus (Pullarkat *et al*., *Phys. Rev. Lett.*, **96**, p.048104, (2006)). If *E* is reduced due to “melting” of the bulk material, the tube can develop peristaltic modes under very low membrane tension. In axons, depolymerization of microtubules generate a fluid-like layer between the stable core and the outer membrane which makes the membrane unstable to beading-like modes. If the core has a thickness gradient, the beads will propagate as in the case of conical wires wetted by a viscous fluid (Lorenceau E & Quéré D, *J. Fluid. Mech.*, **510**(510): p.29–45, (2004)). In axons, the instability can be very periodic when the driving force is strong (high membrane tension) as in osmotic-shock experiments (here only axons which were not attached to the substrate were used). When driving is weak (low membrane tension as in Noco. experiments), structural heterogeneities and adhesive contacts with the growth substrate cause significant dispersion in bead distribution (**Fig. S1**).

## Appendix 3 Bead shapes at long times after Noco treatment

To further test the capillary (i.e., membrane tension) origin of Noco-induced beading, we analyzed the shape of an isolated bead at long times after treatment; see **Fig. S9A**. **Figure 2** shows a remaining microtubule track long after Noco treatment is applied, suggesting a resemblance between axon beading and Rayleigh-Plateau instability-driven dewetting of a liquid film on a cylindrical fiber. In this case, it is well known that the instability saturates to a final pattern of static, isolated droplets. The shape of these droplets can be derived from Laplace’s law: *H* = *P*/(2*σ*), where *H* is the surface mean curvature, *P* is the fluid pressure and *σ* the liquid-vapor interfacial tension. At equilibrium *P* is uniform, and therefore *H* is, as well. From this, the shape of axisymmetric droplets on the fiber can be obtained (B. J. Carroll, *J. Colloid and Interface Science*, **57** p. 488 (1976)).

To see whether axons can be analyzed in a similar manner, we first mathematically characterized the surface of a typical axon bead by determining the radial distance, *R*(*z*), from the bead centerline to the bead contour in the image plane as a function of distance along the axon, *z*; see **Fig. S9A**. Then, by fitting the contour data (**Fig. S9A**) and assuming that the bead is axisymmetric, we calculated the surface principal curvature in the orthoradial direction, 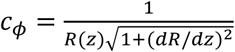 (**Fig. S9B**); the principal curvature in the (*R, z*) plane, 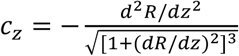 (**Fig. S9C**); and the mean curvature *H* = (*c*_*ϕ*_ + *c*_*z*_)/2, (**Fig. S9D**). Interestingly, there is a central region of the bead where *H* is approximately constant, suggesting that the bead is liquid and with a surface shape governed by a competition between internal pressure and membrane tension. However, the mean curvature increases by roughly a factor of two near the bead edges, where the axon surface is expected to interact with the remaining microtubule core, a behavior which does not occur for liquid droplets wetting a fibre.

We argue that this difference is likely due to the bending rigidity, *κ*, of the membrane enclosing the axon. That is, the shape of the axon surface, *R*(*z*), is determined by a balance of forces *and* torques acting on each surface element; in contrast, the shape of a liquid droplet only involves forces. This translates into requiring greater specification of boundary conditions along the contact line (CL) between the surface and the central fibre than in the liquid case. Namely, *H* is *imposed* along the CL, and is likely quite different than the value given by *P*/(2*σ*). We therefore expect there to be a region of thickness of order 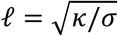 (the characteristic length scale of the membrane), over which *H* varies from the boundary value to the bulk value *P*/(2*σ*). At distances away from the bead edges large compared with ℓ, bending becomes irrelevant and the bead shape is controlled by pressure and tension. Taking *κ* = 3 × 10^−19^ J, as measured for chicken DRG neuronal membrane (Hochmuth *et al.*, Biophys. J., **70**(1), p.358—369 (1996)), and *σ* = 8 × 10^−6^ N/m obtained for our Noco-treated axons, we find ℓ ≈ 200 nm. Note that this value could be modified by the rigidity conferred by the actin-spectrin skeleton. Thus, at long times after axon beading begins, the deformed shape can be understood as arising from a simple balance of forces and torques involving uniform pressure, surface tension, and bending rigidity.

